# Genetic mechanisms of resistance to targeted KRAS inhibition

**DOI:** 10.1101/2025.08.04.668444

**Authors:** Bianca J. Diaz, Max Kops, Sara Bernardo, Henri Schmidt, Elizabeth Grankowsky, Adrian Vega, Chen Zhang, Matthew Bott, Maria Skamagki, Aidan Tomlinson, Nicole A. Vita, Alyna Katti, Mark P. Labrecque, Ida Aronchik, Mallika Singh, Lukas E. Dow

**Affiliations:** Sandra and Edward Meyer Cancer Center, Department of Medicine, Weill Cornell Medicine, New York, NY; Weill Cornell Graduate School of Medical Sciences, Weill Cornell Medicine, New York, NY; Department of Computer Science, Princeton University, Princeton, NJ; Department of Pathology and Laboratory Medicine, Weill Cornell Medicine, New York, NY; Department of Surgery, Memorial Sloan Kettering Cancer Center, New York, NY; Revolution Medicines, Inc., Redwood City, CA; Department of Medicine, Weill Cornell Medicine, New York, NY; Department of Biochemistry, Weill Cornell Medicine, New York, NY

## Abstract

*KRAS* mutations are among the most prevalent oncogenic drivers in non-small cell lung cancer (NSCLC), yet the mechanisms of therapeutic resistance to KRAS inhibitors in these cancers remains poorly understood. Here, we deploy high-throughput CRISPR base editing screens to systematically map resistance mutations to three mechanistically distinct KRAS-targeted therapies, including KRAS-G12C(OFF) inhibitor (adagrasib), RAS(ON) G12C-selective tri-complex inhibitor (RMC-4998), and RAS(ON) multi-selective tri-complex inhibitor (RMC-7977). Using both a saturation *Kras* tiling approach and cancer-associated mutation library, we identify common and compound-selective second-site resistance mutations in *Kras*, as well as gain-of-function and loss-of-function variants across cancer-associated genes that rewire signaling networks in a context-dependent manner. Notably, we identify a recurrent missense mutation in capicua (*Cic*), that promotes resistance to RMC-7977 in vitro and in vivo. Moreover, we show that targeting NFκB signaling in CIC-mutant cells can resensitize them to RAS pathway inhibition and overcome resistance.

## INTRODUCTION

Non-small cell lung cancer (NSCLC) is the leading cause of cancer related deaths worldwide with 5-year survival rate below 20%^1^. Gain-of-function mutations in *KRAS* are the most frequent oncogenic alterations in NSCLCs and recent FDA approval of targeted KRAS-G12C(OFF) inhibitors sotorasib and adagrasib promise a safer and more targeted treatment strategy than cytotoxic chemotherapy^2–5^. In addition to these inhibitors that covalently bind the mutant cysteine in the GDP-bound (inactive; OFF) state, recent work described the development of inhibitors such as RMC-4998 and RMC-7977 that target the GTP-bound (active, ON) state of KRAS. These ‘tri-complex’ inhibitors target GTP-bound G12C-mutant RAS (RMC-4998) or both mutant and wildtype GTP-bound RAS proteins (RMC-7977) via interaction with cyclophilin A (CYPA)^6–8^.

While both adagrasib and sotorasib have shown efficacy in primary and metastatic disease, drug resistance remains a significant problem. Cancer-associated single nucleotide variants (SNVs) have been directly linked to upfront (KEAP1) and acquired (KRAS, MEK etc.) KRAS inhibitor resistance^9–14^. While these examples reveal the potential contribution of SNVs to drug response, the functional impact of the majority of cancer-associated SNVs are poorly understood. CRISPR base editing (BE) technologies enable the targeted creation of SNVs and have been used effectively to explore the genetic mechanisms of drug resistance^15–17^. Indeed, we and others have demonstrated the utility of high throughput BE screens to functionally interrogate variants of unknown significance found in cancers^18–23^. Here, we use BE gene tiling to identify second site mutations in KRAS that drive resistance to three independent RAS-targeted agents with different mechanisms of action. Further, we profile a collection of recurrent cancer-associated mutations, revealing a range of gain and loss-of-function variants to allow escape from one or more RAS inhibitors. We show that a common hotspot mutation in capicua (*CIC*) promotes resistance to the RAS(ON) multi-selective inhibitor RMC-7977 in vitro and in vivo and highlight possible combination therapy routes to overcome resistance.

## RESULTS

To define the spectrum of SNVs in KRAS and other genes that enable drug resistance we chose to use an engineered mouse model that would provide a ‘clean’ genetic background and allow us to isolate the impact of de novo BE-induced mutations. For this, we produced lung tumor-bearing LSL-*Kras*^*G12C*^;*p53*^*fl/ fl*^ (KP) mice following intratracheal delivery of adenoviral-Cre (Figure 1a)^24^ and generated multiple independent KRAS^G12C^ / p53^−/−^ (hereafter “KCP”) tumor cell lines. From these, KCP4 and KCP5 showed the most efficient editing with a GFP BE reporter (Figure 1b)^25^.

**Figure 1.**
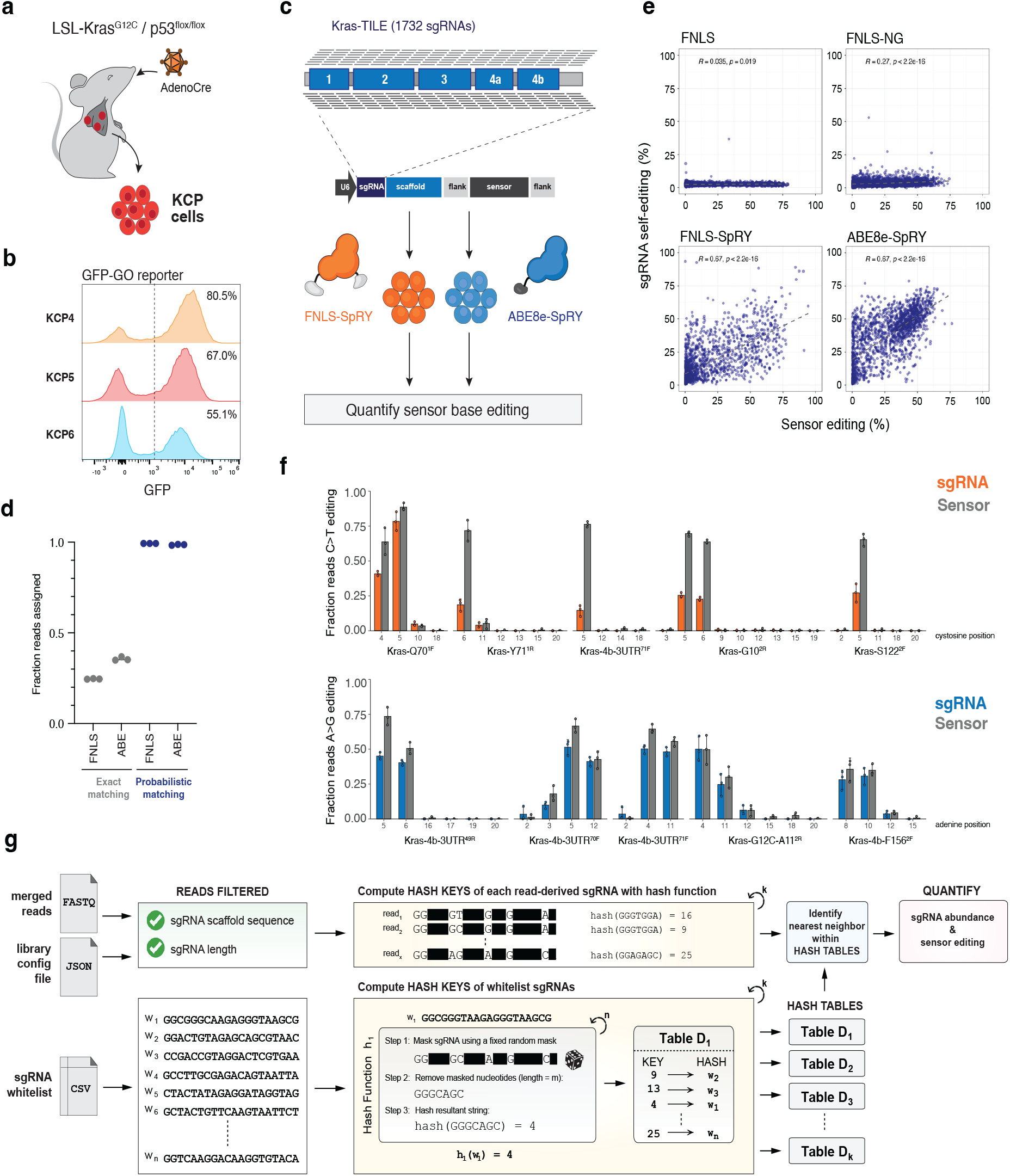
A new computational pipeline to analyze BE tiling libraries. a, Schematic methodology to derive 2D lung tumor lines. b, Measurement of cytosine base editing efficiency with GO reporter by GFP expression using flow cytometry. c, Schematic of the *Kras*-TILE library cloned into our library sensor vector and transduced into cells expressing CBE or ABE. d, Comparison of fraction of reads output from merged fastqs from D0 samples of CBE and ABE cells (n = 3) transduced with *Kras*-TILE either using exact matching or probabilistic matching to the whitelist. e, Correlation of BE editing activity detected at sensor to sgRNA self-editing at D0 across cells transduced with FNLS (NGG), FNLS-NG, and FNLS-SpRY and ABE8e-SpRY. f, Reads were pulled from individual sgRNAs with high BE activity in the *Kras*-TILE library and analyzed with CRISPResso2 at the sgRNA and sensor regions. C>T (orange) and A>G (blue) transitions were measured across all potential nucleotides. g, Schematic describing BEquant pipeline.

We previously described a BE library ‘sensor’ design that enables the quantitation of both sgRNA abundance and an estimation of target editing efficiency at a surrogate target cassette^18^. Using this strategy, we designed a BE Kras saturation or ‘tiling’ library (*Kras*-TILE) with the goal of delineating second-site mutations in *Kras* that would drive resistance to one or more targeted small molecule inhibitors (Supp. Table 1). To maximize the likelihood of identifying most, if not all, possible C>T or A>G resistance mutations, we designed a library to cover every nucleotide position (forward and reverse) across the *Kras* coding sequence, including specific guides to target the G12C-specific mutations present in the L*SL-Kras*^*G12C*^ allele, as well as adjacent intronic and untranslated regions (UTRs); in total, the library consists of 1732 sgRNAs (Figure 1c).

### BEquant: a computational pipeline to analyze BE sensor screens with PAM flexible enzymes

To first test the editing coverage of the library, we transduced KCP4 cells expressing FNLS-SpRY or ABE8e-SpRY at greater than 1000x representation and cultured cells for one week before measuring editing at the sensor (Figure 1c)^26^. Surprisingly, and in contrast to our previous experience^18^, we noted that in both FNLS-SpRY (CBE) and ABE8e-SpRY (ABE) cells, less than a third of filtered reads could be assigned to a sgRNA in the whitelist (Figure 1d). Recently, Ryu et al. reported that PAM-flexible SpRY-based base editors are prone to ‘self-editing,’ whereby the sgRNA itself is modified by the editor^27^. Indeed, while analysis of prior FNLS and FNLS-NG screen data showed minimal sgRNA self-editing, we observed a strong correlation (R = 0.67-0.69) between editing at the sensor and sgRNA sites for both FNLS-SpRY and ABE8e-SpRY enzymes (Figure 1e, Supp. Fig 1a). Further, analysis of editing at individual sgRNA examples from the Kras-TILE libraries reveal similar patterns of C>T or A>G conversion within the sgRNA and sensor editing windows (Figure 1f, Supp. Fig 1b). In the absence of an independent unique molecular identifier (UMI) linked to each sgRNA in the library, self-editing interferes with conventional read assignment workflows that rely on exact matching of each read’s sgRNA to a predefined whitelist. For example, our previous analysis pipeline required exact matches between each read and the whitelist sgRNA and scaffold sequence, excluding any read containing a single mismatch.

To overcome this limitation, we developed a probabilistic matching algorithm based on a locality-sensitive hashing (LSH) workflow popularized in computer science^28^. The logic of the algorithm is described in detail in the Supplementary Methods, but briefly, the approach consists of three steps (Figure 1g). First, reads are preprocessed to remove those containing indels or mismatches in the sgRNA scaffold by performing an exact string match; scaffold mutations are associated with reduced sgRNA editing efficiency (Supp. Fig 2). Second, each 20-nucleotide sgRNA sequence in the whitelist is indexed in a hash table defined by a unique set of randomly selected positions (default: m = 7) (Figure 1g). Specifically, for each sgRNA in the whitelist, a hash key is generated by concatenating the nucleotides at the defined positions and then applying a standard hash function to the concatenated string. The corresponding guide ID is then stored under this hash key. Finally, we compute the hash key of each read-derived sgRNA using the preceding hashing procedure; to identify the best match, the hamming distance (number of mismatches) between the read-derived sgRNA and the whitelist sgRNAs are computed, and the minimal distance whitelist sgRNA is assigned. To improve the probability of finding the nearest whitelist sgRNA to a given read-derived sgRNA, we store multiple hash tables (default: k = 20 tables) each defined by a different set of (m) randomly selected positions (Figure 1g). Once the read-derived sgRNA is assigned to a specific whitelist sgRNA (“guide ID”), quantifying editing outcomes becomes straightforward: count the number and type of base edits at the target site for a specified nucleotide (Figure 1g).

Although one could iterate through all possible whitelist sgRNAs to assign a whitelist sgRNA to each read-derived sgRNA, this approach is computationally prohibitive. The probabilistic matching algorithm reduces the time complexity over this naive procedure from O(n×m) to approximately O(n + m), where n is the number of reads and m is the size of the whitelist. For example, for 1000× read coverage of the Kras-TILE library (n ≈ 1.7 million and m ≈ 1,700), this results in an estimated 1,000-fold improvement in processing time. Most importantly, as compared to our previous exact matching approach, a real-world implementation of the probabilistic matcher increased the percentage of successfully assigned reads from ~25–35% to over 99% in FNLS-SpRY and ABE8e-SpRY datasets (Figure 1d).

### BE tiling libraries identify second site resistance mutations to KRAS inhibitors in vitro

Analysis of sensor editing across the Kras-TILE library showed consistent sgRNA activity across most of the Kras coding sequence, lower overall editing at targets with PAMs containing 2 or more pyrimidines or polyT-rich regions, as expected (Figure 2a, Supp. Fig 3a and b). To identify mutations conferring resistance to KRAS blockade, we treated three independently transduced replicates with either adagrasib (G12C(OFF)), RMC-4998 (RAS(ON) G12C-selective) or RMC-7977 (RAS(ON) multi-selective). Cells were treated with increasing inhibitor concentrations over 30 days in culture and gDNA was collected at days 0, 10, 20 and 30, marking each dose transition (Figure 2b). Libraries from each timepoint were sequenced and sgRNA abundance and sensor editing were measured with BEquant (Supp. Table 3). Enrichment or depletion of individual sgRNAs was quantified using MaGECK (Supp. Table 4)^29^.

**Figure 2.**
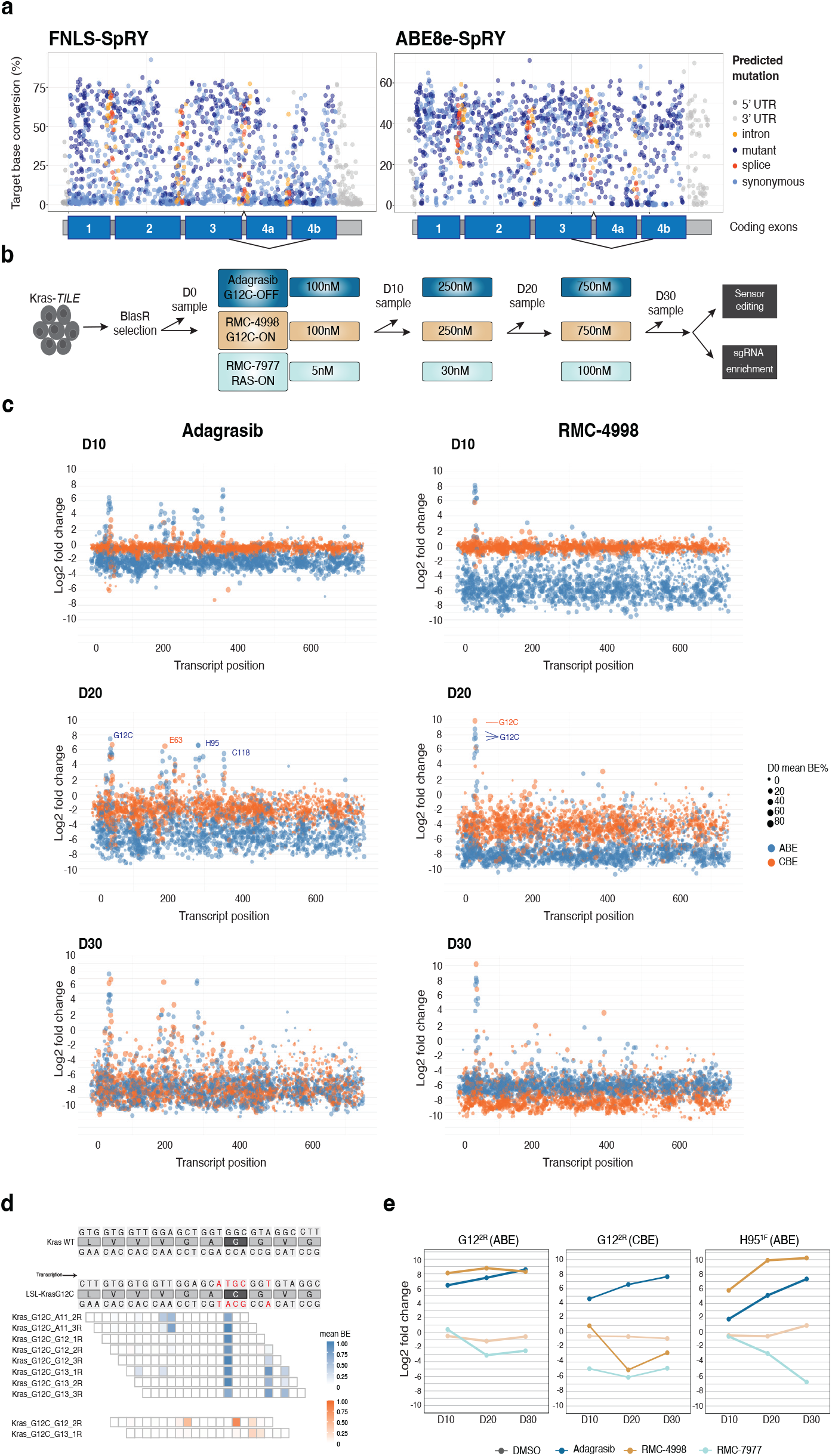
KRAS BE tiling identifies unique and shared hotspot resistance mutations to adagrasib and RMC-4998. a, Percent BE at the target base (cytosine or adenine in the 5th position of sgRNA) across the *Kras* transcript (x axis). Dots are colored by the potential outcome of the predicted mutation. b, Schematic of *Kras*-TILE screen describing dosing and samples for NGS (n = 3 transduction replicates). c, Manhattan plot demonstrating the median log2 fold change of each sgRNA calculated with MAGeCK (y axis) across the Kras transcript (x axis) at each timepoint across Adagrasib and RMC-4998 treated samples (n = 3). d, Schematic of *Kras* WT and LSL-*Kras*^*G12C*^ sequence depicting *Kras*-TILE sgRNAs from Adagrasib and RMC-4998 treated samples. e, Line graphs demonstrating changes in log2 fold change calculated with MAGeCK (y axis) of the G122R and H951F sgRNAs over time (x axis). Lines are colored by drug treatment.

In each treatment setting, enrichment of sgRNAs in both CBE and ABE screens were localized to specific regions of the *Kras* coding sequence. For adagrasib, minimal sgRNA enrichment was observed at day 10 (100nM), but by D20 (250nM) and D30 (750nM) of treatment, there was significant over-representation of sgRNAs targeting the mutant *Kras*^*G12C*^ allele (Figure 2c-e, Supp. Fig 4), consistent with the specific covalent targeting of the mutant cysteine. Adagrasib-treated samples also showed enrichment of sgRNAs targeting the switch II (SWII) region of KRAS, as well as H94/H95/Y96 (Figure 2c and e, Supp Fig 4). Mutations in these regions have previously been associated with clinical adagrasib resistance^10^, strongly supporting the utility of BE screens to identify clinically relevant resistance mutations. RMC-4998 treated samples showed a more restricted pattern of resistance mutations, with the only significantly enriched sgRNAs targeting the mutant *Kras*^*G12C*^ allele (Figure 2c-e and Supp. Fig 4), consistent with covalent engagement of KRAS^G12C^ by the CYPA:RMC-4998 complex. Unlike adagrasib and RMC-4998, RMC-7977 treated samples did not enrich for sgRNAs targeting G12C, consistent with the ability of this small molecule to target multiple KRAS mutants as well as wildtype KRAS, NRAS, and HRAS^6^. Guides targeting the E62/E63/Y64 and Y71 loci were enriched in all treatment doses (Figure 3a and b and Supp. Fig 4). Sequencing of exon 3 in the endogenous *Kras* locus confirmed the strong enrichment of E63K and Y71H mutations, matching the predicted mutations by these sgRNAs (Figure 3c).

**Figure 3.**
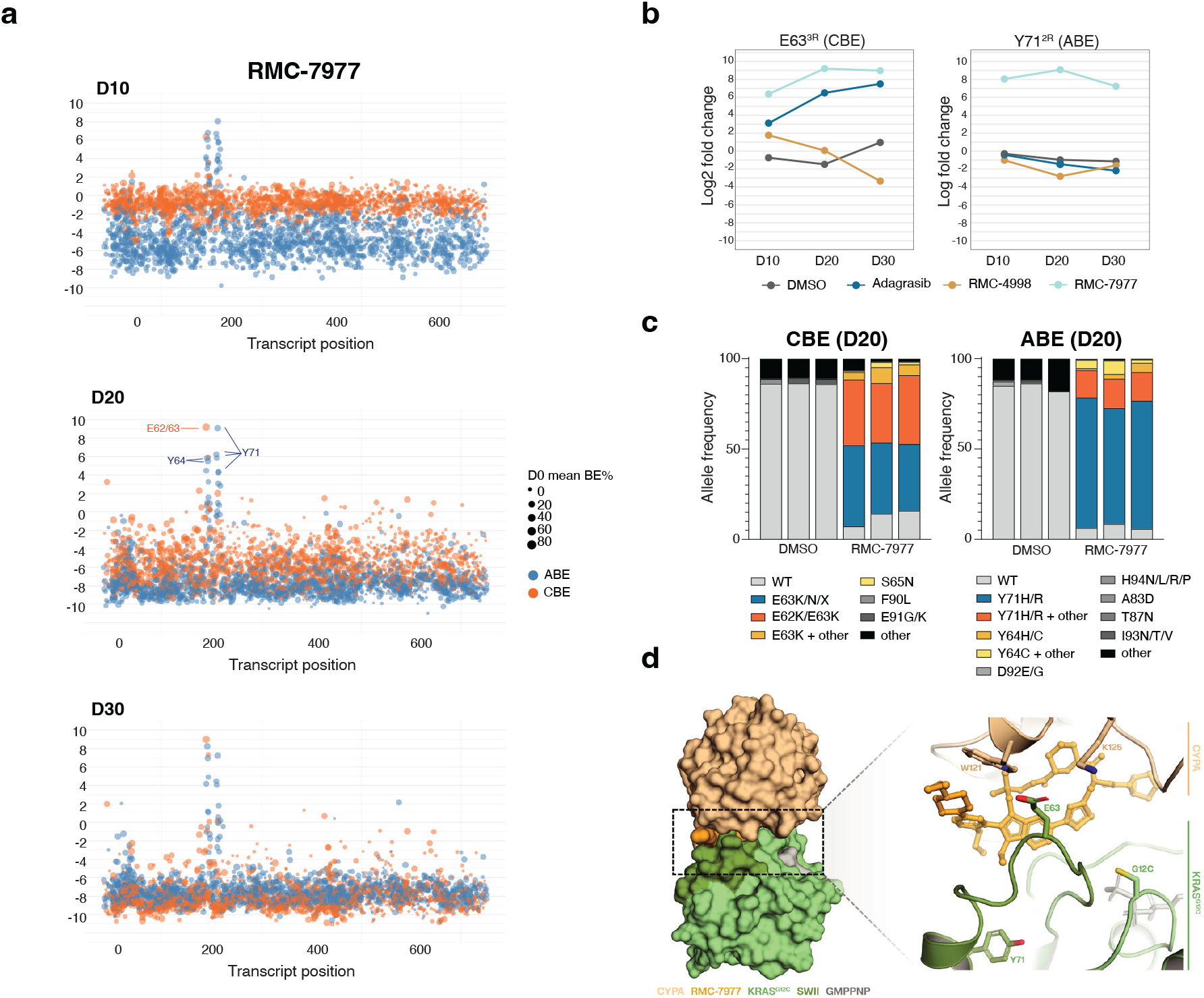
KRAS BE tiling identifies unique and shared hotspot resistance mutations to RMC-7977. **a**. Manhattan plot demonstrating the median log2 fold change of each sgRNA calculated with MAGeCK (y axis) across the Kras transcript (x axis) at each timepoint in RMC-7977 treated samples (n = 3). **b**. Line graphs demonstrating changes in log2 fold change with MAGeCK (y axis) of the E633R and Y712R sgRNAs over time (x axis). Lines are colored by drug treatment. **c**. Stacked bar chart showing the frequency of specific alleles determined by amplification of the exon and NGS of the endogenous locus of Kras from the screen samples treated with DMSO or RMC-7977 at D20 for CBE (left n= 3) and ABE (right n=3). **d**. X-ray crystal structure of CYPA:RMC-7977:KRAS(GMPPNP)G12C (PDB:8TBK) shows that the SWII region of KRAS forms part of the protein-protein interface of the tri-complex. E63 is proximal to both RMC-7977 and CYPA residues W121 and K125 while Y71 forms a number of hydrophobic interactions within KRAS.

The selective enrichment of Y71 and E63 mutations in RMC-7977-treated cells relative to other inhibitors, prompting us to assess the impact of these variants on inhibitor-target interactions. Visual inspection of tri-complex crystal structures show that E63 is at the CYPA-KRAS protein-protein interface, and Y71 folds into a hydrophobic core in the distal SWII region. (Figure 3d and Supp. Fig 5a). We modelled the effect of introducing an additional mutation of E63K or Y71H on tri-complex formation using computational models of RMC-7977 bound to CYPA and KRAS(GMPPNP)G12C/E63K or KRAS(GMPPNP)G12C/Y71H (Supp. Fig 5b and c). Modelling shows that the substitution of E63 to lysine creates both steric and electrostatic clashes with the CYPA residues W121 and K125. Mutating the lipophilic Y71 to a polar histidine leads to increased mobility of the hydrophobic core under SWII with a markedly different conformation of SWII. (Supp. Fig 5b and c) Molecular dynamics simulations of both double mutants show an overall destabilization of the SWII and the protein-protein interface that would lead to reduced complex formation (Supp. Fig 5d). Interestingly, a third resistance variant identified in our screen, Y64X, was recently identified in NSCLC and CRCs that had progressed on treatment with the investigational agent daraxonrasib (RMC-6236), the clinical analog of RMC-7977^30^. Similar to our modeling described above, Y64C/D/H mutations interfered with formation of the daraxonrasib-induced KRAS-CYPA ternary complex^30^.

Together, these data confirm validity of BE sensor screens as an approach to identify genetic drivers of drug resistance and predict second-site KRAS mutations that may drive resistance to the RAS(ON) multi-selective inhibitor RMC-7977.

### A subset of cancer associated mutations drive resistance to KRAS inhibition in KRASG12C NSCLC preclinical models

In addition to acquired second-site mutations in KRAS, other, non-KRAS genetic changes may influence the response to targeted KRAS inhibition. For instance, missense mutations in MAP2K1 and BRAF, among others, have been linked to KRAS inhibitor resistance^9–11^. To define the spectrum of cancer-associated mutations that limit the response to KRAS blockade, we used our previously described mouse BE sensor (MBES) library that contains 4686 sgRNAs designed to create ~1177 recurrent cancer-associated mutations (Supp. Table 1)^18^. This library is comprised of both predicted gain-of-function and loss-of-function mutations, as well as variants of unknown significance (VUS) (Figure 4e). As described for the *Kras*-TILE screen above, FNLS-SpRY expressing KCP4 cells were transduced at 1000X representation followed by antibiotic selection for 7 days (Figure 4a). To improve the statistical power of these screens and limit the identification of false positives due to ‘jackpot’ enrichment in a single replicate, we performed the screen with 5-6 replicates for each treatment arm. Further, since we noted minimal difference in the identification of hits between day 20 and day 30 in the *Kras*-TILE screen, we restricted our MBES screen to D20 (Figure 3a) (Supp. Tables 5 and 6).

**Figure 4.**
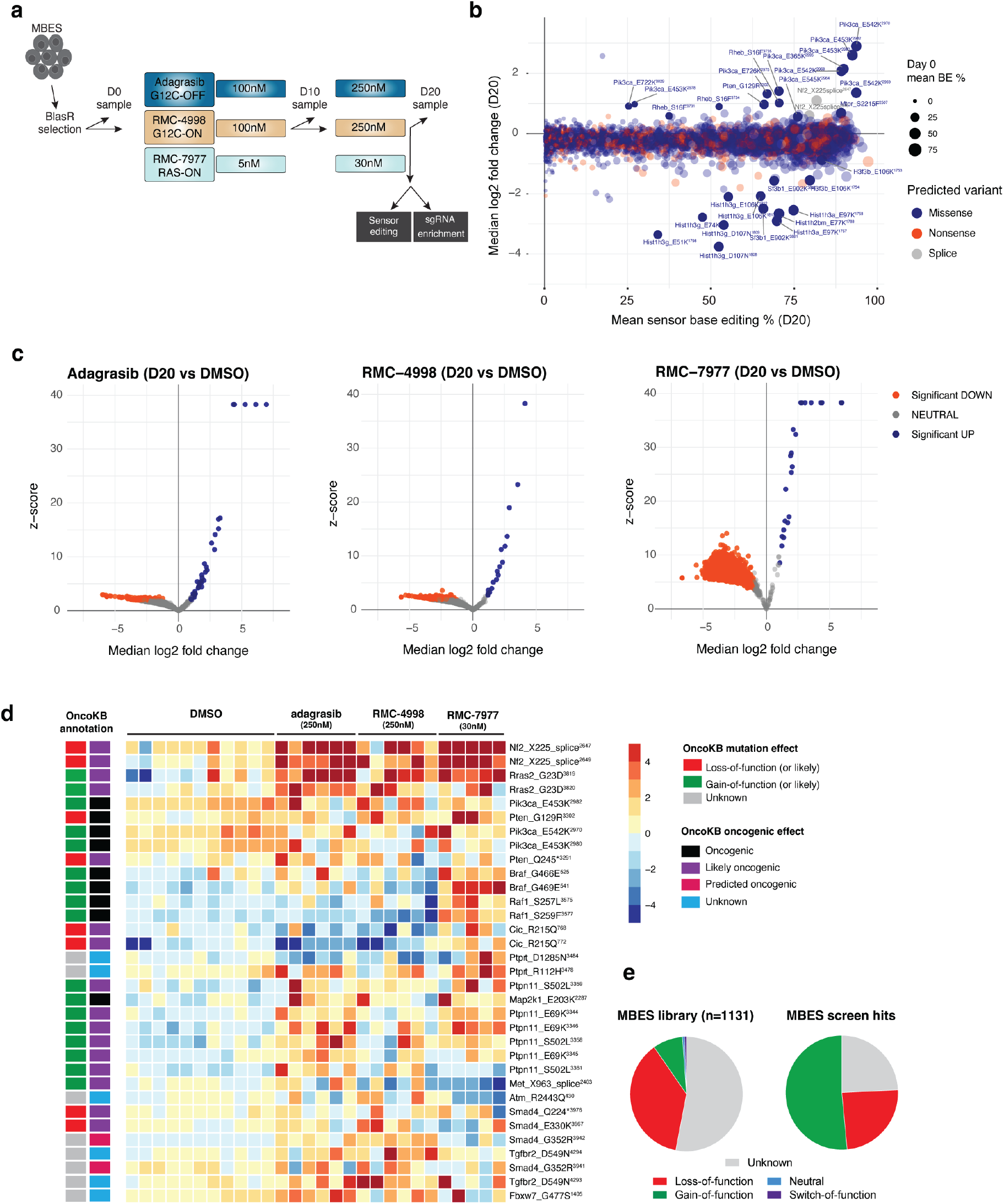
MAPK and inflammatory regulators drive resistance to KRAS inhibitors. a, MBES screen timeline schematic describing dosing and samples for NGS (n = 5-6 transduction replicates). b, Bubbleplot comparing sgRNA log2 fold change (y axis) and mean sensor base editing (x axis) at D20 in DMSO control treated samples. Colors represent predicted variant and size of dot represents mean BE% at D0. c, Volcano plots of sgRNAs that are significantly enriched in blue and significantly depleted in orange from Adagrasib, RMC-4998, and RMC-7977 treated samples compared to DMSO at D20 (left to right). d, (left) OncoKB annotation of each enriched sgRNA at D20 and its predicted mutation outcome and its predicted oncogenic effect. (right) Heatmap of D20 enriched sgRNAs as rows and each column representing a replicate and binned by its associated treatment condition. Each box is colored by its log2 fold change score calculated with MAGeCK. e, Pie charts (left) of predicted functional outcomes of sgRNAs in the MBES library and (right) of predicted functional outcomes of sgRNAs from the enriched hits at D20.

Analysis of untreated (DMSO) samples after 20 days of culture highlighted a collection of enriched sgRNAs targeting regulators of PI3K/MTOR signaling, including those targeting *Pik3ca, Pten, Rheb, Mtor*, and *Nf2*. There were a similar number of significantly depleted sgRNAs, all representing histone and splicing factor variants (Figure 4b). The reduced abundance of cells carrying these sgRNAs is consistent with the observation that “oncohistone” and splicing factor mutations are almost always identified as heterozygote alleles, whereas homozygous alterations negatively impact cell fitness^31,32^.

All inhibitor treated samples showed a large proportion of neutral or significantly depleted sgRNAs, with a small number of enriched sgRNAs in each inhibitor arm. In total, across all inhibitor treatments we identified 33 sgRNAs representing 23 distinct SNVs, that were significantly enriched (median LFC > 1, p-adj < 0.05) in one or more conditions, and showed editing greater than 20% at the sensor. Interestingly, while only ~11% of the MBES library model predicted gain-of-function mutations, they represented more than half the resistance hits (17/33) (Figure 4c). The hits could be broadly categorized by those common across all inhibitors, only in mutant-selective inhibitors, or only in RMC-7977 treated samples. The strongest hits across all treated samples were sgRNAs targeting an *Nf2*^*x225*^ splice site and sgRNAs targeting *Rras2*^*G23*^ - predicted to be a gain of function mutation akin to G12X variants in other RAS family members (Figure 4d). In addition, we identified multiple sgRNAs targeting regulators of MAPK signaling, including predicted gain and loss-of-function mutations both upstream (*Ptpn11, Met*) and downstream (*Map2k1, Raf1, Braf*) of RAS (Figure 4d). Further, sgRNAs predicted to create activating mutations PI3K signaling were enriched in inhibitor treated samples, consistent with previous clinical data from adagrasib-treated patients^10^. In mutant-selective inhibitor treated samples, multiple sgRNAs targeting *Tgfbr2* and *Smad4* were enriched by Day 20 (Figure 4d), while in RMC-7977 treated samples, we identified multiple sgRNAs predicted to drive gain-of-function mutations in *Raf1* and *Braf*, as well as mutations in the transcriptional repressor capicua (*Cic*)^33^ (Figure 4d).

As a first step to validate the relative resistance induced by some of the top hits, we cloned individual guides targeting *Rras2*^*G23*^, *Nf2*^*x255*^, or *Cic*^*R215*^ into a tdTomato-P2A-BlasR vector and transduced KCP-FNLS-SpRY cells. TdTomato-positive sgRNA-expressing cells were mixed at a 20:80 ratio with GFP-positive KCP-FNLS-SpRY parental control cells and the percentage of each cell population was monitored over time by flow cytometry, following treatment with each inhibitor. We observed a strong enrichment of both the RRAS2^G23^ and CIC^R215^ mutant population across all treatment conditions, while surprisingly, the NF2^x225^ mutant cells showed more modest enrichment (Figure 5a, Supp. Fig 6a). To better assess the impact of each specific mutations across multiple cell models, we generated single cell clones from each mutant in two independent KCP lines (KCP4 and KCP5; n=3 clones/ mutation/line) (Supp. Fig 6b). Interestingly, NF2^x225^ KCP4 clones showed resistance under most inhibitor doses, while KCP5 clones with the same mutations behaved like controls cells (Supp. Figure 7a and b), highlighting significant heterogeneity between these mouse tumor models. RRAS2^G23D/N^ and CIC^R215Q^ KCP4 clones showed some variability in response to inhibitor treatment, while KCP5 clones demonstrated consistent outgrowth under inhibitor treatment (Figure 5b and c and Supp. Fig 7c). Overall, CIC^R215Q^ mutant clones showed the most consistent behavior under inhibitor treatment and represented the only individual mutation tested that enabled significant resistance to RMC-7977 in both KCP lines (Figure 5b and c, Supp. Fig 7c, Supp. Table 9). Cells carrying the RRAS2^G23D/N^ mutation showed attenuated suppression of pERK over 48hrs of drug treatment, consistent with the ability of RMC-7977 to induce tri-complex formation between RRAS2 and CYPA, albeit at reduced efficiency^6^ (Figure 5d). In contrast, CIC^R215Q^ mutant clones showed no changes in pERK or pS6 compared to CR8 control clones (Figure 5d, Supp. Fig 7d). To confirm the impact of CIC disruption was not specific to murine cells, we engineered a CIC^R215Q^ mutations in Calu-1 NSCLC cells that are KRAS^G12C^ / p53 mutant, reflecting the genotype of the mouse tumor model. Consistent with data from mouse cells, CIC^R215Q^ mutant Calu-1 clones showed ~4-fold reduced sensitivity to RMC-7977 in vitro (Figure 5e, Supp. Fig 8, and Supp. Table 9).

**Figure 5.**
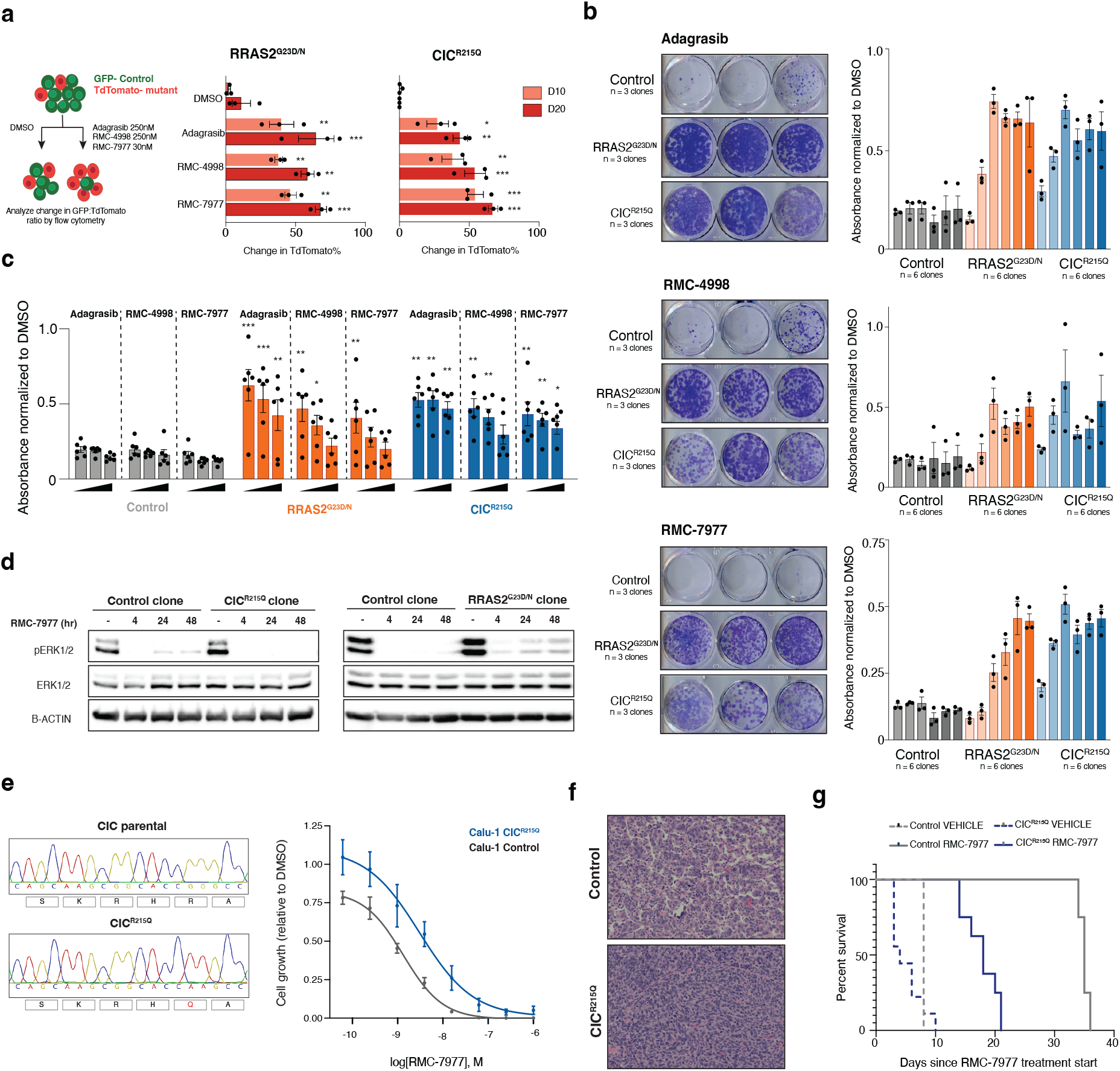
CIC mutations promote resistance to multi-selective RAS inhibition in mouse and human cells and in vivo in NSCLC. a, (left) Schematic of the competition assay methodology. KCP4, 5, and 6 cells are individually transduced with a control sgRNA in a GFP expressing guide plasmid and a mutant sgRNA in a TdTomato expressing guide plasmid and selected for 7 days with blasticidin. Selected cells are mixed in a 80:20 control to mutant cell ratio and either treated with DMSO, Adagrasib (250nM), RMC-4998 (250nM), and RMC-7977 (30nM). Samples were analyzed for the GFP and TdTomato expression at D3 (initial timepoint) and D10 and D20. (right) Quantification of change in positive TdTomato population % from D3 at D10 and D20 from each treatment group in Rras2^G23D/N^ and Cic^R215Q^ mutant cells (n = 3 biological replicates and performed 2 independent times). Two-way ANOVA (analysis of variance); *P < 0.05, **P < 0.01 ***P < 0.0001. b, (left) Representative images of endpoint of colony formation assay of 3 independently derived singlcell clones after treatment with Adagrasib (250nM), RMC-4998 (250nM), and RMC-7977 (30nM). (right) Quantification of Giemsa stain as measured by absorbance normalized to DMSO treated cells for 6 independently derived clones per mutant in KCP4 and KCP5 cells. c, Summary quantification of all colony formation assay experiments performed. Each bar is an increasing dose of Adagrasib (100nM, 250nM, and 750nM), RMC-4998 (100nM, 250nM, and 750nM), and RMC-7977 (5nM, 30nM, and 100nM). Each dot represents the mean absorbance value of an individual single cell clone performed 3 independent times (n = 2 independent cell lines and 3 independent single cell clones). Two-way ANOVA (analysis of variance); *P < 0.05, **P < 0.01 ***P < 0.0001. d, Western blotting of control and mutant single cell clones from KCP4 cells after treatment with RMC-7977 30nM and protein samples collected at 4, 24, and 48 hours where expression of phosphorylated ERK is assessed. e, (left) Sanger sequencing traces of CIC R215 sgRNA locus from Calu-1 control and Calu-1 CIC^R215Q^ single cell clone. (right) RMC-7977 dose response assay comparing cell viability at D6. Data represents 3 independently derived single cell clones and performed 2 separate times. Two-way ANOVA (analysis of variance); ***P < 0.0001. f, Representative H&E images from vehicle treated control or Cic^R15Q^ mice (n = 3-5/group) g, Survival analysis of mice transplanted with control or Cic^R215Q^ cells (n = 3-5/group). **P < 0.01. P values were calculated using the log-rank test.

To test whether CIC^R215Q^ mutations drive therapy failure in vivo, we engrafted CR8 control or CIC^R215Q^ mutant KCP5 cells into the lung by tail vein injection and began daily RMC-7977 treatment (10 mg/kg) after 2 weeks. To avoid potential issues with immune rejection from the expression of Cas9 in KCP5 clones, we used inducible BE transgenic mice (iBE) as hosts^34^. CR8 control KCP5 cells appeared as moderately well differentiated adenocarcinomas maintaining some glandular and/or alveolar architecture, while CIC mutant tumors were sarcomatoid, spindly, and pleomorphic, with a solid growth pattern completely effacing the underlying alveolar architecture (Figure 5f). As expected, there was no significant survival difference between vehicle-treated CR8 and CIC mutant transplanted animals (Figure 5g). Treatment with RMC-7977 induced prolonged survival benefit in mice carrying CR8 control tumors, showing an increased survival benefit of 27 days compared to vehicle treated animals (Figure 5g, p=0.008). CIC mutant tumors showed partial response to RMC-7977, but had significant, 50% reduction in median survival relative to drug-treated CR8 tumors (27d vs 14d, p=0.003) (Supp. Table 9). Together, these data support the results from the MBES screen and confirm that CIC^R215Q^ mutations promote resistance to mutant-specific and multi-selective RAS inhibitors in multiple lung cancer models with KRAS-G12C oncogenic mutations.

### Identifying targetable dependencies induced in CIC^R215Q^ mutant cells

CIC has previously been characterized as a transcriptional repressor of MAPK and YAP signaling and its loss has been linked to MEK inhibitor resistance in human cancer cell lines^33,35–39^. To explore the potential mechanism(s) underlying RMC-7977 resistance in KRAS-G12C models harboring CIC^R215Q^ mutations, we performed RNA-seq on KCP4 and KCP5 control and mutant lines following acute treatment (D3) and following prolonged treatment (D20), after all control cells had been eliminated. Gene enrichment analysis revealed upregulation of various transcriptional programs across both control and CIC^R215Q^ mutant cells, including epithelial-to-mesenchymal transition (EMT), YAP signaling, and both AT1 and AT2 cell identity signatures (Supp. Fig 9 and Supp. Tables 7 and 8). Interestingly, despite a known role as a MAPK transcriptional repressor, only relatively few genes in the KRAS signaling gene set showed clear derepression in CIC mutant cells, including *Etv4* and *Etv5*, that have previously been linked to CIC-dependent MEK inhibitor resistance (Figure 6a, upper panel)^38^. In contrast, roughly half of previously published CIC targets^35,39^ were derepressed in CIC^R215Q^ mutant cells (Figure 6a, middle panel). We focused our attention on programs that were significantly altered in CIC^R215Q^ cells, but not in control, NF2^x225^ or RRAS2^G23D/N^ mutant cells. This analysis revealed a small subset of transcriptional programs, of which only NFκB signaling showed positive regulation in CIC^R215Q^ mutant cells and negative enrichment scores in CR8 controls (Figure 6b). Interestingly, over-representation analysis identified KRAS, ER, and NFκB as gene sets enriched in previously described CIC genomic targets (Figure 6c). Further, several NFκB target genes were upregulated in RMC-7977 treated CIC^R215Q^ mutant cells, but largely unchanged in control clones (Figure 6a, lower panel). To determine whether targeting NFκB signaling would block resistant outgrowth in CIC^R215Q^ mutant cells, we co-treated control and CIC^R215Q^-mutant cells with RMC-7977 and the small molecule inhibitor IKK-16 that selectively targets Iκb kinase (IKK-1, IKK-2, and IKK complex). While CR8 control cells showed minimal impact of IKK-16 treatment and no enhanced effect of the inhibitor combination, IKK-16 synergized with RMC-7977 to block proliferation of CIC^R215Q^ mutant cells. (Figure 6d-f). Moreover, pre-treated RMC-7977-resistant cells showed enhanced growth arrest under combination treatment (Figure 6d and Supp. Table 9). These findings suggest that CIC^R215Q^-mediated resistance to RMC-7977 is driven, at least in part, by inflammation-associated transcriptional changes, and targeting the NFκB pathway offers a potential strategy to overcome this resistance and restore sensitivity to RAS(ON) multi-selective inhibition.

**Figure 6:**
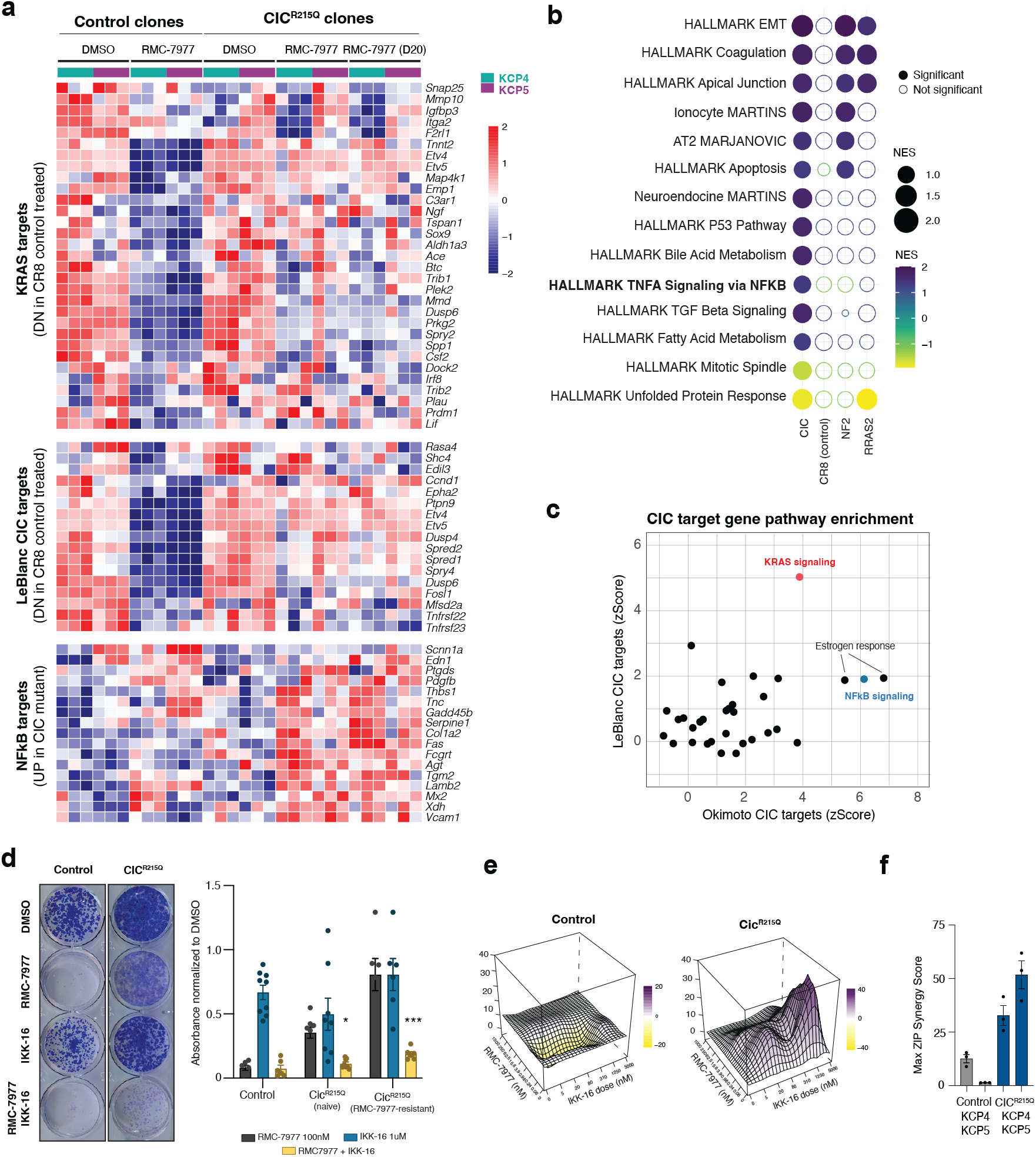
Targeting NFκB synergizes with RMC-7977 to suppress outgrowth of CIC mutant cells. a, Heatmap showing average Log2 normalized expression of differentially expressed genes from KRAS targets, LeBlanc CIC targets, and NFkB targets gene sets across DMSO and RMC-7977 30nm treated samples (n = 6 independently derived single cell clones from KCP4 and KCP5 cells). b, Summary of the gene sets significantly enriched in CIC^R215Q^ mutant samples compared to control, NF2^x225_splice^, and RRAS2^G23D/N^ mutant samples treated with RMC-7977 30nM. Size and color of dot represents NES. Shading of dot represents significance (n = 6 independently derived single cell clones from KCP4 and KCP5 cells). c, Dotplot comparing CIC target pathway enrichment z scores from two published datasets Okimoto CIC targets (x axis) and LeBlanc CIC targets (y axis). d, (left) Representative image of colony formation assay experiment of endpoint of control and CIC^R215Q^ cells treated with RMC-7977 30nM, IKK-16 1uM, or the combination. (right) Quantification of Giemsa stain absorbance value normalized to DMSO. (n = 3 independently derived single cell clones from KCP5 cells and performed 2 separate times). Two-way ANOVA (analysis of variance); *P < 0.05 and ***P < 0.0001. e, Control and CIC^R215Q^ mutant cells were treated with indicated concentration of DMSO, RMC-7977, and IKK-16 for 6 days and synergy was calculated based on cell viability at different dose pairs. Data are mean of 3 biological replicates. f, Summary of Max synergy scores across control and CIC^R215Q^ mutant samples.

## DISCUSSION

The advent of KRAS targeted therapies in NSCLC have provided meaningful clinical benefits, however, resistance to these treatments is common. Our study leverages CRISPR BE screens to systematically interrogate the genetic landscape of KRAS inhibitor resistance in NSCLC. Using high-throughput BE sensor screens, we identified clinically validated and previously unidentified second-site mutations in KRAS that drive resistance to KRAS G12C(OFF) and RAS(ON) G12C- and multi-selective inhibitors. Consistent with early clinical data, mutations in codons H95 and Y96 emerged as resistance hotspots unique to adagrasib^10,40^, highlighting the utility of BE screens in identifying relevant resistance variants. Interestingly, two small molecules inhibitors that bind GTP-bound KRAS through a tri-complex interaction with CYPA showed distinct resistance mutation profiles. RMC-7977-treated samples enriched strongly for mutations in E63, Y64, S65, and Y71 residues located in the SWII region, along the interface between KRAS and CYPA. Consistent with an important role for these residues in target inhibition, recent clinical analysis identified Y64C/D as a driver of resistance to daraxonrasib through disruption of tri-complex formation^30^. In contrast, RMC-4998, despite binding KRAS through the same tri-complex mechanism, showed exquisite selectivity for sgRNAs targeting the G12C mutation. These data reveal key potential modes of resistance to mutant-selective and multi-selective inhibitors in ongoing clinical trials, and importantly, suggest that resistance to one inhibitor may not preclude sensitivity to a secondary treatment. Mapping the cross-sensitivity of new and existing small molecule KRAS inhibitors using *Kras*-TILE (or similar approaches) could enable effective second-line treatment strategies for tumors that develop second-site resistance variants.

Unlike many other BE tiling screens that have previously been reported^17,19–23,41^, our study exploited SpRY-based BE variants, enabling true saturation tiling of the coding sequence, with minimal PAM bias. The consequence of ‘near-PAMless’ sequence recognition was significant self-editing at the sgRNA itself. This phenomenon was also recently reported by Ryu et al^27^ and interferes with the enumeration of sgRNAs through standard sequence-matching approaches. To overcome this problem, we developed a new computational pipeline, BEquant, that uses locality sensitive hashing and nearest neighbor matching to provide efficient assignment or edited or non-edited sgRNAs without the need for additional unique molecular identifiers (UMIs)^42^. Thus, the BEquant pipeline is a computationally efficient and flexible approach for analyzing sensor-based library screens, and importantly, empowers the use of near-PAMless editing enzymes for full coverage saturation BE screens.

In addition to KRAS second-site mutations, we identified a collection of cancer-associated mutations that can bypass KRAS inhibition, highlighting multiple potential pre-existing or acquired alterations that could limit the extent and/or durability of therapy. The majority of hits were regulators of MAPK signaling, consistent with strong dependence of KRAS/MAPK signaling in KRAS mutant tumors. Of note, the identification of multiple class 3 Braf mutations (G466E and G469E) that are in part RAS-dependent and induce lower overall pERK activation^43^ suggests that even moderate MAPK activation may drive drug resistance^30^. We also identified multiple mutations in genes involved in inflammatory signaling, including *Tgfbr2, Smad4*, and *Ptprt*^44–46^, implying a role for cell intrinsic inflammatory signals in KRAS inhibitor resistance.

One mutation that enabled robust RMC-7977 resistance in multiple KRAS-G12C NSCLC tumor models in vitro and in vivo was a variant in the DNA binding region of *Cic*. Beyond the known role of CIC in repression of MAPK and YAP transcriptional targets, we identified a specific deregulation of NFκB signaling targets. Interestingly, CIC has been shown to directly bind the Drosophila NFκB homolog, Dorsal47, and more recently, has been shown to constrain aberrant expression of interferon stimulated genes (ISGs) in mammalian cells^48,49^. The induction of a subset of ISGs in CIC mutants identified NFκB signaling as a potential vulnerability, and treatment with the NFκB inhibitor IKK-16 effectively restored sensitivity to RMC-7977. These findings align with previous studies implicating NFκB signaling in oncogenic adaptation^50^ and are consistent with our recent description of non-genetic mechanisms of bypass to KRAS/EGFR dual therapy in colorectal cancer. These studies raise the possibility that in some cases, genetic and non-genetic pathways co-opt similar mechanisms to evade treatment.

Overall, our study provides a framework for identifying and characterizing resistance mutations in cancer models treated with targeted therapies. Our findings support the predominant role of MAPK pathway reactivation as a mechanism of resistance to diverse KRAS inhibitors and in select cases, identify cell intrinsic inflammatory signaling as a potential therapeutic vulnerability in KRAS-G12C NSCLC. The recognition that CIC can regulate both MAPK and inflammatory targets raises the question of whether other MAPK and/or inflammatory regulators identified in the screen, such as TGFBR2 and PTPRT converge on a similar mechanism that could be targeted to improve the outcome of KRAS targeted therapy.

## Acknowledgements

We would like to acknowledge members of the Dow lab for advice and assistance and the WCM genomics core facility for assistance with the Illumina sequencing of sgRNA and transcriptomic libraries. We thank John Knox for thoughtful review and constructive comments. This work was supported by a Sponsored Research Agreement from Revolution Medicines and an Emerald Distinguished Investigator Award to LED. BJD was supported by an F31 award from the National Cancer Institute (NIH/NCI) under award number (F31-CA261061-01). AV was supported by a Human Frontiers Postdoctoral Fellowship. LED is the Burt Gwirtzman Research Scholar in Lung Cancer at Weill Cornell Medicine

## Author Contributions

BJD, MK, SB designed and performed experiments, analyzed data, generated figure panels and wrote the paper. HS developed the computational pipeline, designed figures and wrote the paper. EG, AV, MB, MS, AT and NAV performed experiments. AK, MPL, IA, MS performed analysis and supervised the work. LED designed experiments, analyzed data, supervised the work and wrote the paper.

## Data Availability

Sensor amplicon sequencing and RNA sequencing raw fastq files are available at the sequence read archive (SRA) under accession: PRJNA1293657

## METHODS

### Plasmids and individual sgRNA cloning

The following lentiviral base editing plasmids were used in this manuscript: FNLS-SpRY and ABE8e-SpRY, (this manuscript), all new plasmids and libraries will be made available on Addgene. The following sgRNA plasmids were used in this manuscript: LRT2Bv2 and LRG2Bv2 (this manuscript). We cloned BsmBI-compatible annealed and phosphorylated oligos encoding sgRNAs into BsmBI-linearized LRT2Bv2 using high concentration T4 DNA ligase (NEB). A 5’ G (to boost U6 transcriptional initiation) was added to sgRNAs that lacked it either by appending it to the 5’ or by substituting the first nucleotide in the 5’ position for a G. All sgRNA sequences used are listed in Supplementary Table 2.

### Design and construction of *Kras*-TILE library

#### Base editing sensor module design

Each sensor module is composed of the following parts: 1) a 22nt long 5’ adapter/priming site with a Esp3I restriction site; 2) a 20nt long 5’ G-containing sgRNA; 3) a 93nt long improved SpCas9 sgRNA scaffold partially based on7; 4) an 11nt long sequence corresponding to the 5’ flanking sequence of the endogenous target site; 5) the 23nt cognate target site; 6) a 7nt long sequence corresponding to the 3’ flanking sequence of the endogenous target site; 7) and a 28nt long 3’ adapter/priming site with a EcoRI restriction site. Thus, oligos encoding individual sensor modules are 204nt long.

#### Design and cloning of mKras tiling library

sgRNAs were designed to target the coding sequence of the mouse Kras sequence along with around ~60 bp of intronic sequence to capture splice site mutations and spaced 1 bp apart on both the positive and negative strands. mKras tiling library oligos were synthesized with using Twist Biosciences. Libraries were cloned into the LRT2Bv2 backbone as follows (all library oligos are in Supplementary Table 1). Briefly, each oligo pool was amplified using forward and reverse primers that append Esp3I and EcoRI sites to the 5’ and 3’ ends of the sensor insert, purified using the QIAquick PCR Purification Kit (Qiagen), and ligated into Esp3I-digested and dephosphorylated pLRT2B vector using high-concentration T4 DNA ligase (NEB) (all cloning and sequencing oligos are in Supplementary Table 1). To ensure maximum library recovery, we set up n=24 parallel PCR reactions per pool. A minimum of

2.4 ug of ligated LRT2Bv2 plasmid DNA per pool (corresponding to n=8 ligations) was electroporated into Endura electrocompetent cells (Lucigen), recovered for one hour at 37C, plated across four 15cm LB-Carbenicillin plates (Teknova), and incubated at 37°C for 16 hours. The total number of bacterial colonies per pool was quantified using serial dilution plates to ensure a library representation of >10,000X. The next morning, bacterial colonies were scraped and briefly expanded for 4 hours at 37°C in 500mL of LB-Carbenicillin. Plasmid DNA was isolated using the Plasmid Plus Maxi Kit (Qiagen). To assess sensor distribution and fidelity of assembly per pool, we amplified the sensor region using primers that append Illumina sequencing adapters on the 5’ and 3’ ends of the amplicon, as well as a random nucleotide stagger and unique demultiplexing barcode on the 5’ end (Supplementary Table 1). Library amplicons were size-selected on a 2% agarose gel, purified using the QIAquick Gel Extraction Kit (Qiagen), and sequenced on an Illumina MiSeq instrument.

### Cell culture

HEK293T (ATCC CRL-3216), KCP4 (derived by Dr. Matthew Bott), KCP5 (derived by Dr. Matthew Bott), and KCP6 (derived by Dr. Matthew Bott), were cultured in DMEM supplemented with 10% fetal bovine serum (FBS) and 100 IU/mL of penicillin/streptomycin. Calu-1 cells were a kind gift of Dr. John Ferrarone (Weill Cornell) and were cultured in RPMI supplemented with 10% fetal bovine serum (FBS) and 100 IU/mL of penicillin/streptomycin.

### Virus production

Lentiviruses were produced by co-transfection of HEK293T cells with the relevant lentiviral transfer vector and packaging vectors psPax2 (Addgene, #12260) and pMD2.G (Addgene, #12259) using PEI MAX (Polysciences). Viral supernatants were collected at 48 and 72 hours post transfection and stored at −80°C.

### Drug treatments

Adagrasib (MRTX849, MedChemExpress Cat. No.: HY-130149) was dissolved in DMSO at a stock concentration of 10 mM and used at final concentrations of 100nM, 250nM, and 750nM. RMC-4998 and RMC-7977 were provided under collaborative agreement with Revolution Medicines, Inc. and were dissolved in DMSO at a stock concentration of 10mM and used at final concentrations of 100nM, 250nM, and 750nM and 5nM, 30nM, and 100nM respectively. IKK-16 (Selleckchem Cat. No. S2882) and was dissolved in DMSO at a stock concentration of 10 mM and used at final concentrations of 1uM.

### Flow cytometric analyses

GO validation and competition assay experiments were measured in a Thermo Fisher 2018 Attune NxT flow cytometer^18^.

#### Analysis of base editing activity using GO reporter system

Base editor expressing cells were plated at a density of 5,000 cells/well in 12 well plates and transduced 24 hours later with a defined amount of GO reporter to achieve 20-50% transduction efficiency. Virus-containing media was replaced with complete media 24 hours post-transduction and cells were harvested for flow cytometry at 96 hours post-transduction. We used an Attune NxT flow cytometer (Thermo Fisher). Cells were trypsinized with a 100 μl of 0.25% Trypsin+EDTA and resuspended in 300 μl of complete medium in a 96 well U bottom plate. Data was acquired at a flow rate of 500 μl/min and at least 10,000 events from the single cell population gating were recorded.

### Screening of Kras-TILE sensor library

We first transduced KCP4, 5, 6 cells with FNLS-SpRY and tested BE activity as detailed above and determine to continue screens with KCP4 cells as they had highest BE activity. KCP4 CBE and ABE cells were transduced with a volume of viral supernatant that would achieve an MOI between ~0.3-0.5. For titering of library virus for standard transduction, cells were plated at densities that mimicked a 1000-2000X library representation (depending on the size of the library) and transduced 24 hours later with increasing volumes of viral supernatant (0, 25, 100, 200, 500, 1000, and 2000 μL) and polybrene (10 μg/mL, EMD Millipore). Cells were incubated at 37°C for 72 hours after which viral infection efficiency was determined by the percentage of tdTomato+ cells assessed by flow cytometry on an Attune NxT flow cytometer. *Kras*-TILE screens were performed in triplicate and each step of the screen – from infection to sequencing – was optimized to achieve a minimum representation of over 2000X. 24 hours after infection, cells from each corresponding replicate were pooled into a minimum of 2 × 150mm tissue culture dishes (Corning) and selected with Blasticidin S (Gibco) at an empirically-determined final concentration of 5 μg/mL. Cells were cultured and kept under Blasticidin selection for seven days post-transduction. When needed, cells were trypsinized and re-plated at the cell number to ensure a minimum representation of 2000X. Subsequently, a D0 sample/replicate was pelleted and stored at −20°C. Cells were split into DMSO, adagrasib (100nM), RMC-4998 (100nM), and RMC-7977 (5nM) and cultured for 10 days at this dose splitting when necessary and ensuring representation a D10 sample/replicate was collected. Next, Cells were split into DMSO, adagrasib (250nM), RMC-4998 (250nM), and RMC-7977 (30nM) and cultured for another 10 days at this dose splitting when necessary and ensuring representation a D20 sample/replicate was collected. Lastly, cells were split into DMSO, adagrasib (750nM), RMC-4998 750nM), and RMC-7977 100nM) and cultured for another 10 days at this dose splitting when necessary and ensuring representation a D30 sample/replicate was collected. Raw BEquant and MAGeCK results in Supplementary table 3 and 4 respectively.

### Screening of MBES library

Methodology is essentially the same as with the Kras-TILE library with the following changes: adagrasib and RMC-4998 arms (n = 6 replicates) and RMC-7977 (n = 5 replicates), Screen cells were only cultured in drug until D20. Raw BEquant and MAGeCK results in Supplementary Table 5 and 6 respectively.

### Deconvolution of Kras-TILE and MBES screens

We assumed that each cell contains approximately 6.6 pg of gDNA. Therefore, screen deconvolution at 1000X required sampling ~6 million x 6.6 pg of gDNA, or ~39.6 ug and performed Genomic DNA extractions using the protocol found on dowlab.org/protocols. We employed a modified 2-step PCR version of the protocol published by Doench et al adapted to our unique library design. Briefly, we performed an initial “enrichment” PCR, whereby the integrated sensor cassettes were amplified from gDNA, followed by a second PCR to append Illumina sequencing adapters on the 5’ and 3’ ends of the amplicon, as well as a random nucleotide stagger and unique demultiplexing barcode on the 5’ end. Each “PCR1” reaction contained 10μL of Herculase II 5X Master Mix (Agilent), 0.5 uL dNTPs, 2.5 μL of Sensor_ v6_Fwd Primer (10 μM), 2.5μL of Sensor_v6_Rev Primer (10 μM), 1 uL of Herculase II polymerase, and 5 μg of gDNA in 33.5 μL of water (for a total volume of 50 μL per reaction). The number of PCR1 reactions was scaled accordingly; therefore, we performed eight PCR1 reactions per technical replicate and time point for all screens. PCR1 amplicons were purified using the QIAquick PCR Purification Kit (Qiagen) and used as template for “PCR2” reactions. Each PCR2 reaction contained either 25 μL of Taq 2X Master Mix (NEB), 1 μL of a uniquely barcoded PCR2_Fwd Primer (10 μM), 1 μL of a common PCR2_Rev Primer (10 μM), and 300 ng of PCR1 product in 20 μL of water (for a total volume of 50 μL per reaction). We performed two PCR2 reactions per PCR1 product. Library amplicons were size-selected either on a 2% agarose gel and purified using the QIAquick Gel Extraction Kit (Qiagen) and sequenced on an Illumina NoveSeq (150 nt paired-end reads). All primer sequences are available in (Supplementary Table 2). PCR program for PCR1 using Herculase II was: 1) 95°C x 30s; 2) 95°C x 10s; 3) 55°C x 30s; 4) 72°C x 30s; 5) Go to step 2 × 24 cycles; 6) 72°C x 2 min; 7) 4°C forever. PCR program for PCR2 using Taq 2X Master Mix (NEB) was: 1) 95°C x 30s; 2) 95°C x 30s; 3) 60°C x 30s; 4) 68°C x 30s; 5) Go to step 2 × 10 cycles; 6) 72°C x 5 min; 7) 4°C forever.

### Analysis of Kras-TILE and MBES data

To quantify base editing outcomes, raw paired-end FASTQ reads were paired using Pandaseq and merged FASTQ files were used as input for downstream analysis using BEquant (Supplementary methods). To quantify sgRNA enrichment and depletion total read counts for each replicate were used as input for MAGeCK analysis. Any sgRNA with read counts < 100 were removed from analysis. Comparisons of DMSO vs treatment at D10, D20, and D30 for each replicate were performed using MAGeCK to determine median log2 fold changes.

#### Computational modeling of KRAS-G12C-RMC-7977-CYPA tri-complexes

Molecular dynamics was performed using Schrodinger 2025-1 Desmond. Schrodinger was used to generate CYPA:RMC-7977:KRAS(GTP)^G12C/E63K^ and CYPA:RMC-7977:KRAS(GTP)^G12C/Y71H^ double mutants followed by a short minimization. The tricomplex structures were then solvated using SPC water in a rhombic dodecahedron xy-hexagon box with a 15 Å buffer. The systems were then neutralized using sodium counter ions and brought up to a salt concentration of 0.15M NaCl. Total simulation time of 100ns using NPT ensemble at 300K and 1.01bar. The RMSD of RAS switch II (residues 57-75) across each frame of the simulation was measured. The data was exported to GraphPad Prism 10.4.2 to calculate the area under the curve and was graphed using MatPlotLib. The structure images from Figure 3d and Supplementary Figure 5a, b and c were generated using PyMol.

### Validation experiments

#### Individual sgRNA infection and confirmation of base edit

For validation of individual targets, sgRNAs were cloned into the lentiviral guide expression vector LRT2Bv2 or LRG2Bv2 and lentiviral particles were produced as described above. Base editor expressing cells were plated at a density of 25,000 cells/well in 12 well plates and were infected 24-hours later with enough virus to achieve 50% transduction efficiency. Virus-containing media was replaced with complete media 24 hours post-transduction and cells were plated into selection media containing 5 μg/mL Blasticidin S (Gibco). Experimental cells remained in selection media until the final collection time point at 7 days post-transduction. Genomic DNA was isolated using the protocol found on dowlab.org/protocols, and targets were amplified using a 100 μl reaction following the standard NEB Taq 2x MM protocol with primers found in (Supplementary Table 1). Each PCR was performed 2X/ target and pooled. Amplicons were confirmed on a 2% agarose gel and PCR purified using QIAGEN QIAquick PCR purification kit. DNA concentration was measured using a Nanodrop and samples were normalized to 20 ng/μl and sequenced using Sanger sequencing by GENEWIZ, Inc (South Plainfield, NJ, USA).

#### Isolation of single cell clones

Control and mutant cells were sorted using the Symphony S6. Cells were gated on size, single cells, DAPI, and tdTomato+ and distributed into 96 well plates as single cells. Cells were monitored for outgrowth of clones and 24 colonies were analyzed per mutant/per cell line and amplified and sequenced as detailed above.

#### Competition assay

Bulk infected cell lines from each validation mutant expressing tdTomato were mixed with control cells expressing GFP at 80:20 ratio in 12 well plates 40,000 control cells:10,000 mutant cells. The following day cells were placed in DMSO, adagrasib (250nM), RMC-4998 (250nM), and RMC-7977 (30nM) containing media. At D3 in drug GFP and tdTomato expression were assessed by flow cytometry using the Thermo Fisher 2018 Attune NxT, Remaining cells were continually split into drug and samples were analyzed again at D10 and D20.

#### Colony formation assay

Cells were seeded at 1000 cells per well into 12-well plates and incubated for 24h for cells to adhere. Compounds were added to the media and were replenished every three days. Once colonies formed (about 6-7 days after compounds were added), the media was removed, and the cells were fixed in 4% paraformaldehyde for one hour at RT on an orbital shaker. The PFA was removed, and the fixed cells were stained in a 10% Giemsa solution in PBS overnight on an orbital shaker. After staining, the Giemsa solution was removed, the plates were washed in water and left to dry overnight. Once dry, the plates were imaged using an Epson Perfection V550 Photo scanner. To quantify cell growth, 1mL of 10% acetic acid was added to each well. The plate was shaken for 20 minutes, and 100ul of the acetic acid solution was transferred to a 96well plate. Absorbance was quantified using a BMG Labtech Fluostar Omega microplate reader.

### Western blot

2e6 (DMSO and 4HRs), 1e6 (24HRs), 500K (48HRs) cells were seeded in 10 cm plates and incubated for 24 hours for the cells to adhere. Compounds were then added to the cells and incubated for 4, 24, and 48 hours. To isolate protein, cells were washed in PBS and lysed in 200ul RIPA buffer on ice for 20 minutes. Lysates were centrifuged at 15,000 rpm for 10 minutes and the protein supernatant was stored at −80C. 25ug of protein was run per sample for western blot analysis. Antibodies used were B-actin (Abcam ab49900), Vinculin (Cell Signaling #18799), p-Akt (Cell Signaling #4060), p—p44/42 MAPK (Cell Signaling #4370), p-S6 (Cell Signaling #4858), Akt (Cell Signaling #2920), p44/42 MAPK (Cell Signaling #9107), and S6 (Cell Signaling #2317).

### RNA isolation and RNA-seq

One six-well of cells was plated per condition into DMSO or RMC-7977 30nM. Cells were cultured and collected at D3 and D20. The cells were collected in 1mL of TRIzol (Thermo Fisher Scientific, #15596018) and RNA was extracted following the manufacturer’s protocol. DNA was removed from the isolated RNA by treating with DNase1 for 15 minutes followed by column purification with the Qiagen RNeasy kit (Qiagen #74106). RNA sequencing was performed by the Genomics Core Laboratory at Weill Cornell Medicine: the RNA quality was confirmed using a 2100 Bioanalyzer (Agilent technologies), the RNA library was prepared using TruSeq Stranded mRNA Sample Library Preparation Kit (Illumina), and RNA-seq was performed on an Illumina NovaSeq 6000 with paired-end 2×100 cycles.

#### RNA-seq analysis

Transcript abundance was estimated using Kallisto51, aligned to the GRCm38 mouse reference genome. Transcript per million (TPM) data was reported for each gene after mapping gene symbols to ensemble IDs using R packages, “tximport”, tximportData”, “ensembldb”, and “EnsDb. Mmusculus.v79”. Differential gene expression was estimated using DESeq252. For data visualization and gene ranking, log fold changes were adjusted using lfcShrink in DESeq2, to minimize the effect size of poorly expressed genes. GSEA analysis (v3.0) was performed on pre-ranked gene sets from differential expression between control and treated groups. We used R (v3.6.1) and R Studio (v1.2.1335) to create all visualizations, perform hierarchical clustering and principal component analysis. Visualizations of RNAseq data were produced using pheatmap (heatmaps), and ggplot2 (dotplots).

### Synergy

Cells were plated at 200 cells/well in 96 well plates and allowed 24 hours to adhere. The next day RMC-7977 and IKK-16 were added at specified doses and drug media was refreshed on the third day. On D6 cell viability (Promega G9241) was assessed using a BMG Labtech Fluostar Omega microplate reader. Synergy was calculated using the SyngergyFinder 3.0.1 R package.

