## Supplementary Figures for "Genetic mechanisms of resistance to targeted KRAS inhibition"

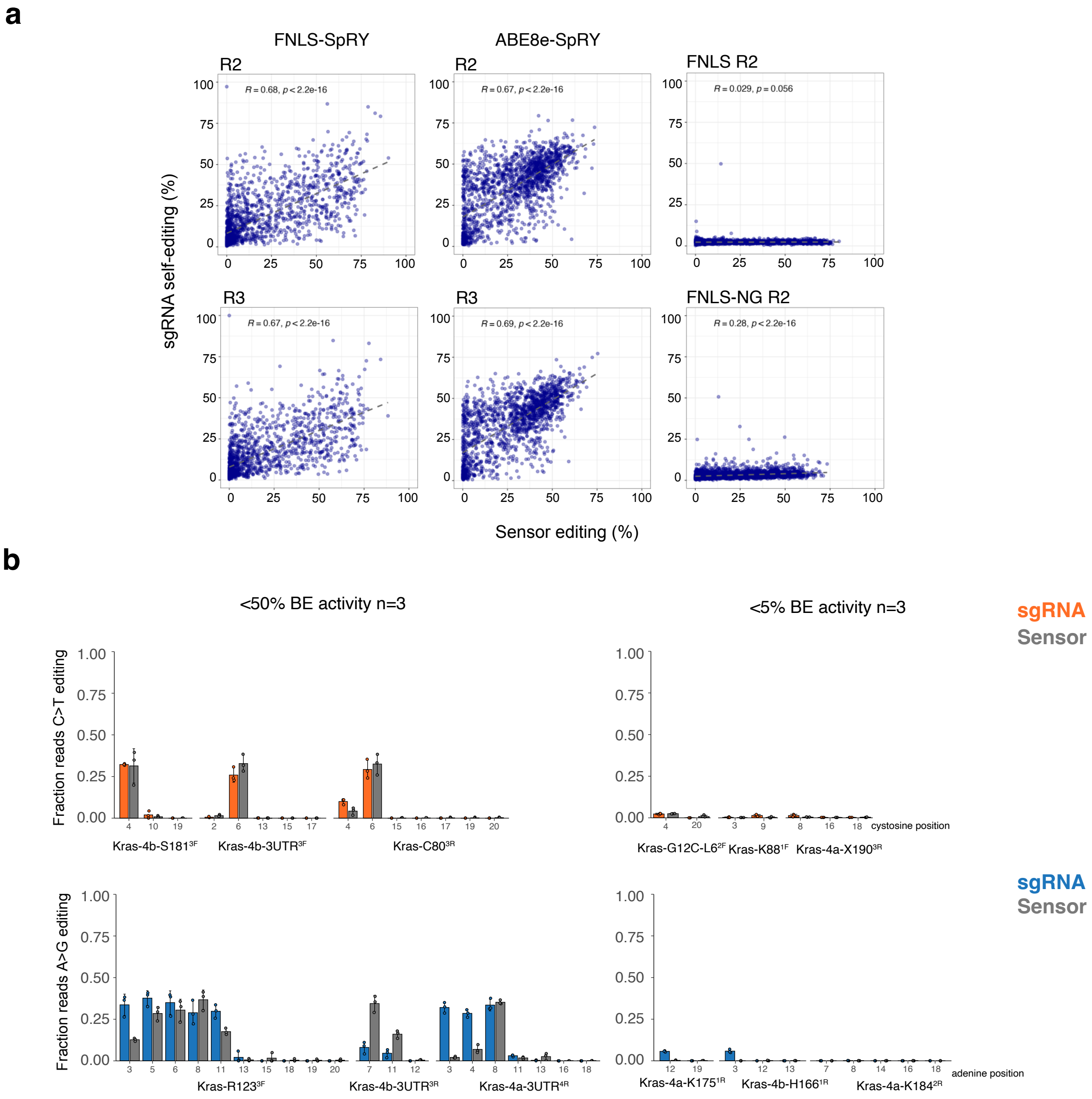


**Supplementary Figure 1. Self-editing of sgRNAs is a feature of SpRY-based editors and follows editing patterns at sensor sites. a,** Correlation of %BE at sensor compared to %BE at sgRNA from D0 samples of FNLS-SpRY and ABE-SpRY cells transduced with *Kras*-TILE library and from FNLS and FNLS-NG cells transduced with MBES. **b,** Reads were pulled from individual sgRNAs with <50% and <5% BE activity in the *Kras*-TILE library D0 samples and analyzed with CRISPResso2 at the sgRNA and sensor regions. C>T (orange) and A>G (blue) transitions were measured across all potential nucleotides (n=3).

**a**

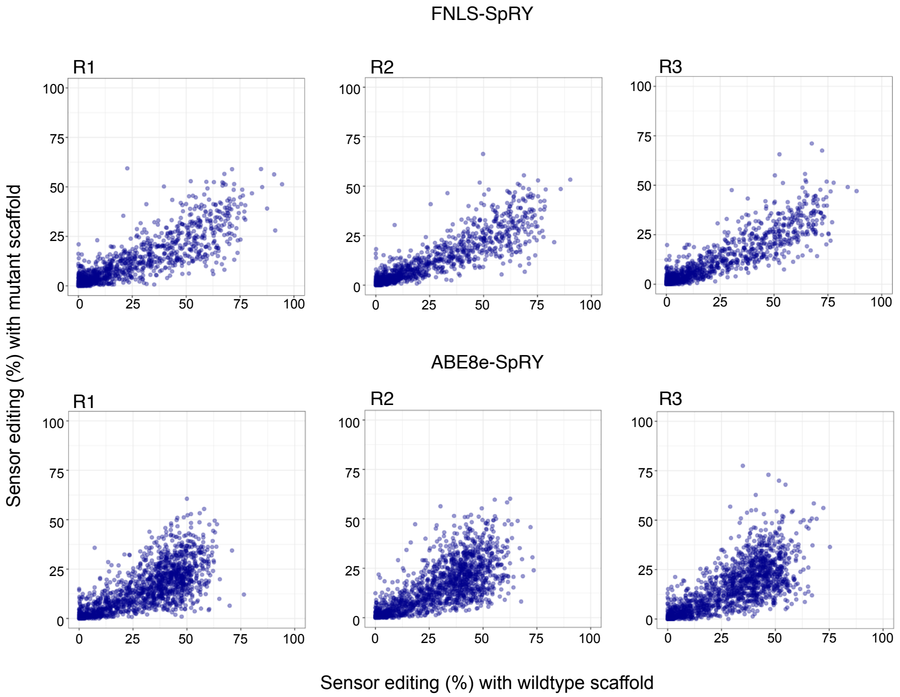


**Supplementary Figure 2. Mutations in the sgRNA scaffold reduce target editing efficiency.** Correlation of D0 %BE at sensor in reads that contain a perfect wildtype sgRNA scaffold and reads with mutant sgRNA scaffold in both FNLS-SpRY and ABE8e-SpRY cells transduced with Kras-TILE library (n = 3).

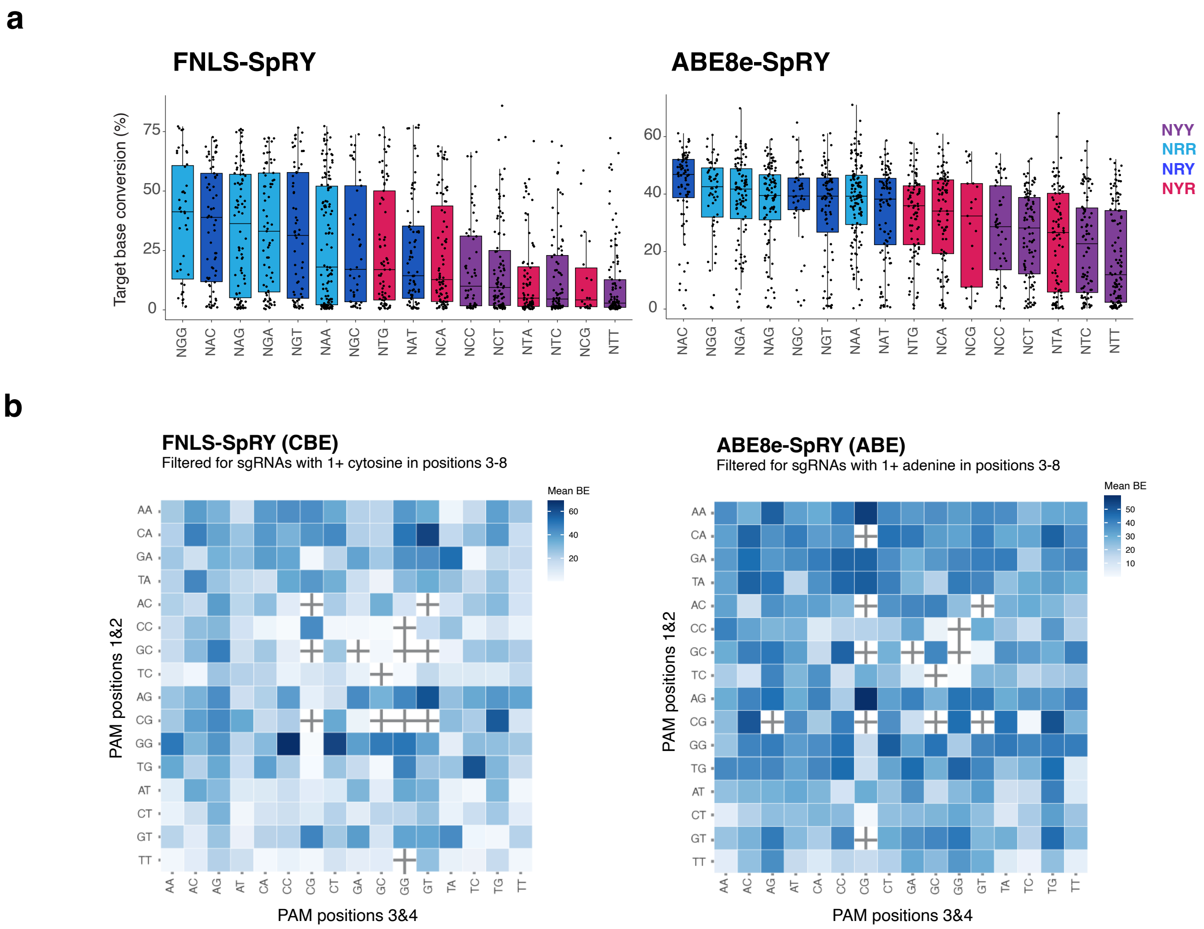


**Supplementary Figure 3. Editing of FNLS-SpRY and ABE8e-SpRY by PAM context and position. a,** Percent CBE (left) or ABE (right) at the target base categorized by the PAM dinucleotide context of the protospacer across sgRNAs from the *Kras*-TILE library. Box plots show the median and interquartile range (IQR) and whiskers represent 1.5× IQR. Dots represent individual data points. **b,** Heatmap of FNLS-SpRY and ABE8e-SpRY BE activity across *Kras*-TILE sgRNAs by PAM positions 1&2 on y axis and PAM positions 3&4 on x axis. Colored by mean BE% and filtered by sgRNAs that contain an adenine or cytosine in the editing window.

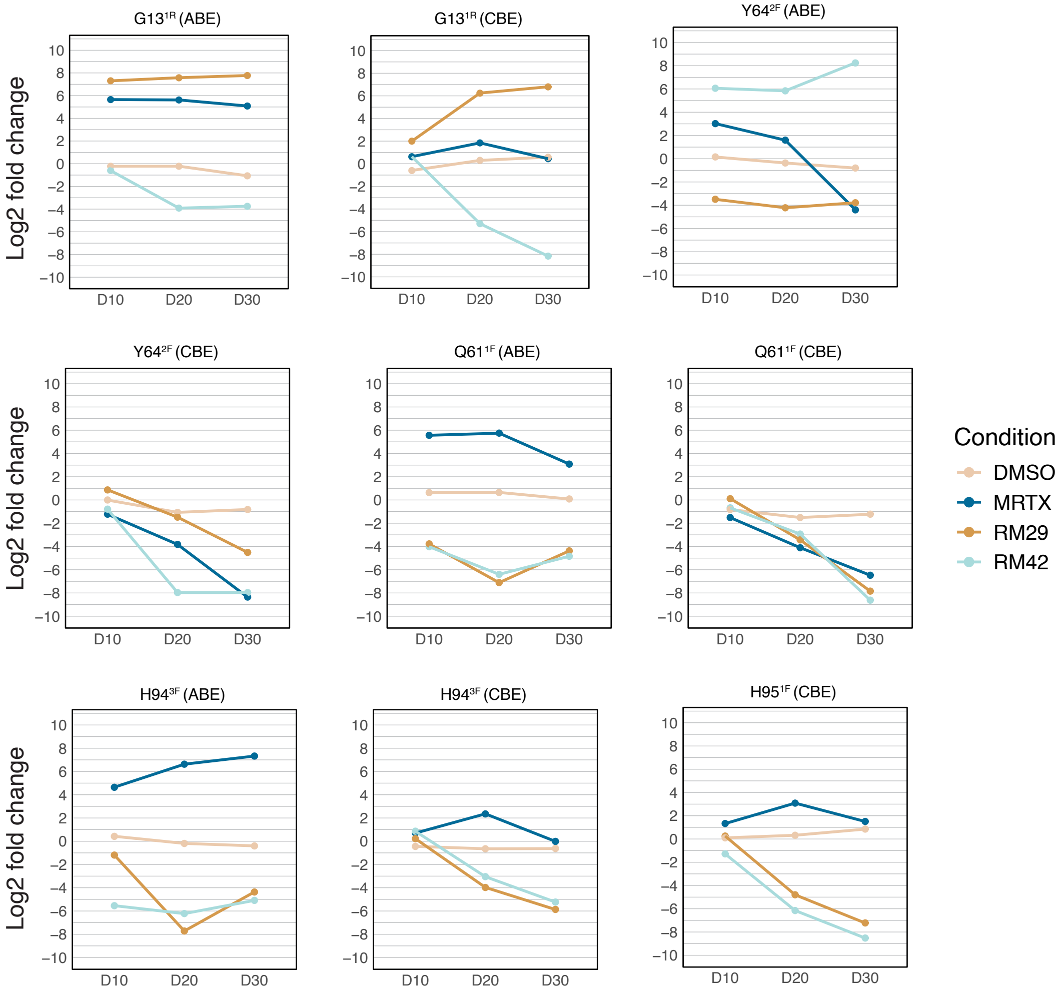


**Supplementary Figure 4. Enrichment and depletion of *Kras*-TILE sgRNAs over time.** Line graphs demonstrating changes in log2 fold change calculated with MAGeCK (y axis) of the *Kras*-TILE sgRNAs over time (x axis). Lines are colored by drug treatment.

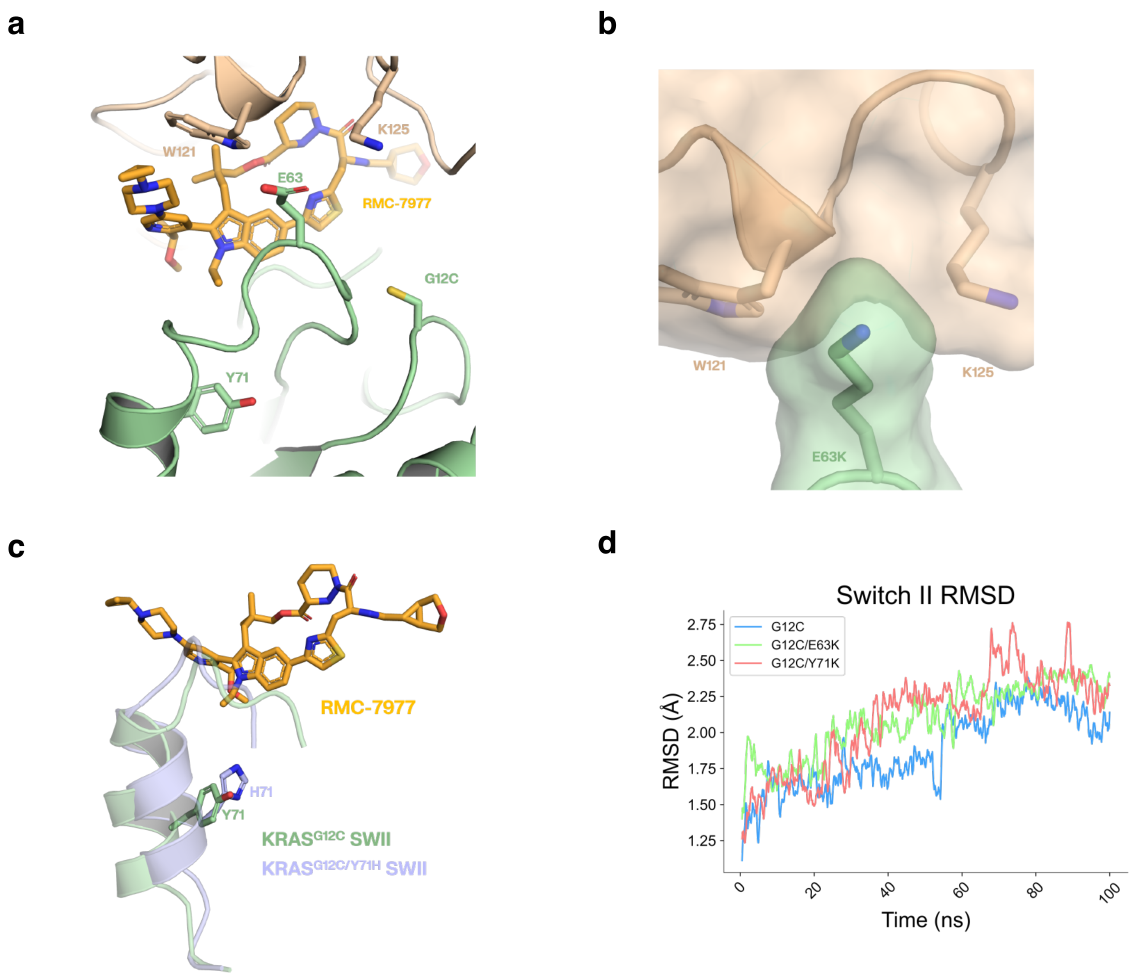


**Supplementary Figure 5. Computational modeling of CYPA:RMC-7977:KRAS(GMPPNP)G12C tri-complexes. a**, E63 and Y71 are located in the SWII region of KRAS, adjacent to the tri-complex interface. **b**, In silico mutation from glutamate to the larger and oppositely charged lysine in the KRASG12C/E63K double mutant suggests potential steric and electrostatic clashes with CYPA interfacial residues W121 and K125, interactions that would disrupt complex formation. **c**, Y71 helps form the hydrophobic core under SWII. **d**, Switch II (residues 57 – 75) RMSD analyses show there is destabilization of the switch II region in both KRASG12C/E63K and KRASG12C/Y71H with larger conformational changes compared to the switch II region of KRASG12C-CYPA-RMC-7977.

**a**

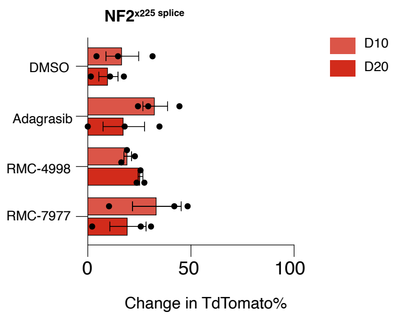


**b**

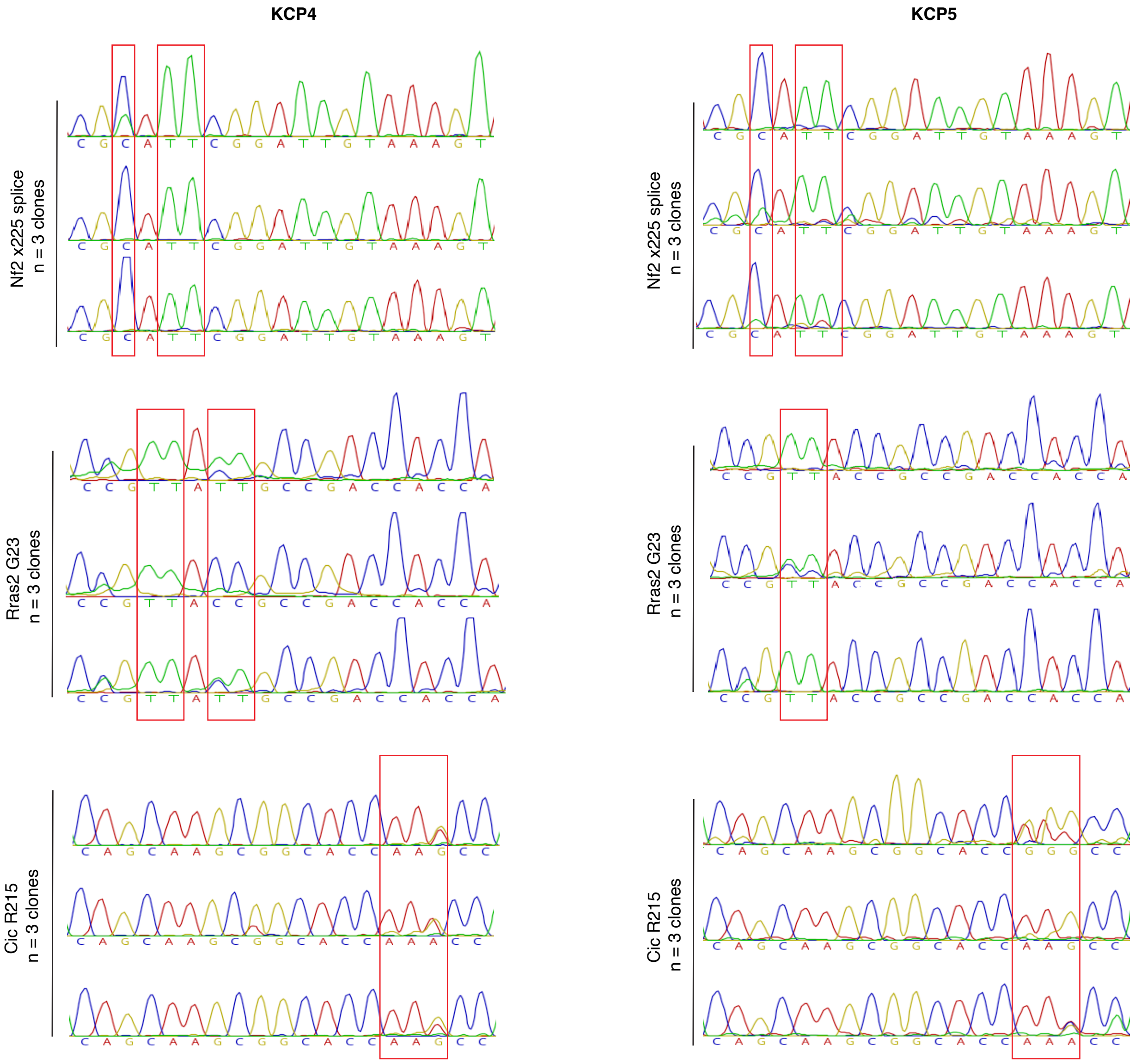


**Supplementary Figure 6. Clone sequencing and western blot characterization. a,** Quantification of competition assay analyzing change in positive TdTomato population % from D3 at D10 and D20 from each treatment group in Nf2$^{x225}$ splice mutant cells (n = 3 biological replicates and performed 2 independent times). **b,** Sanger traces to validate genotype of each single cell clone from Nf2$^{x225}$ splice, Rras2$^{G23D/N}$, and Cic$^{R215Q}$ base edited mutants from KCP4 and KCP5 cells.

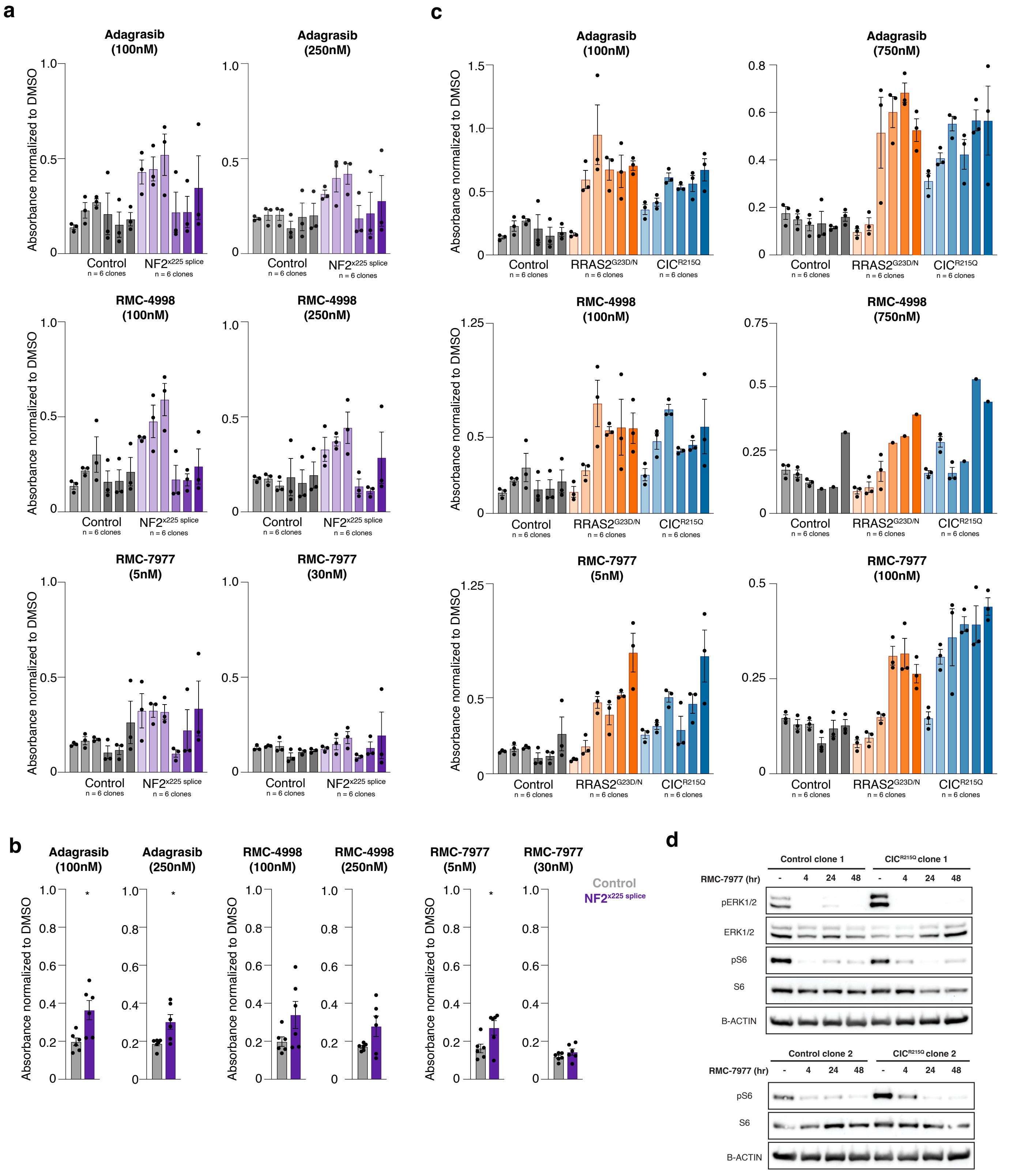


**Supplementary Figure 7. Response of KCP clones to KRAS inhibitor treatment.** **a,** Quantification of Giemsa stain as measured by absorbance normalized to DMSO treated cells for 6 independently derived Nf2$^{x225}$ splice clones per mutant in KCP4 and KCP5 cells treated with Adagrasib (100nM and 250nM), RMC-4998 (100nM and 250nM), and RMC-7977 (5nM and 30nM). **b,** Summary quantification of all colony formation assay experiments performed. Each dot represents the mean absorbance value of an individual single cell clone performed 3 independent times (n = 2 independent cell lines and 3 independent single cell clones). Two-way ANOVA (analysis of variance); $^{*}P < 0.05$. **c,** Quantification of Giemsa stain as measured by absorbance normalized to DMSO treated cells for 6 independently derived Rras2$^{G23D/N}$, and Cic$^{R215Q}$ clones per mutant in KCP4 and KCP5 cells treated with Adagrasib (100nM and 750nM), RMC-4998 (100nM and 750nM), and RMC-7977 (5nM and 100nM). **d,** Western blotting of control and mutant single cell clones from KCP4 cells after treatment with RMC-7977 30nM and protein samples collected at 4, 24, and 48 hours where expression of phosphorylated ERK and phosphorylated S6 is assessed.

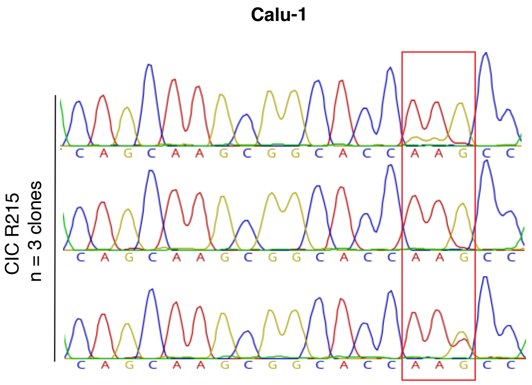


**Supplementary Figure 8. Clone sequencing.** Sanger traces to validate genotype of each single cell clone from $CIC^{R215Q}$ base edited mutants from Calu-1 cells.

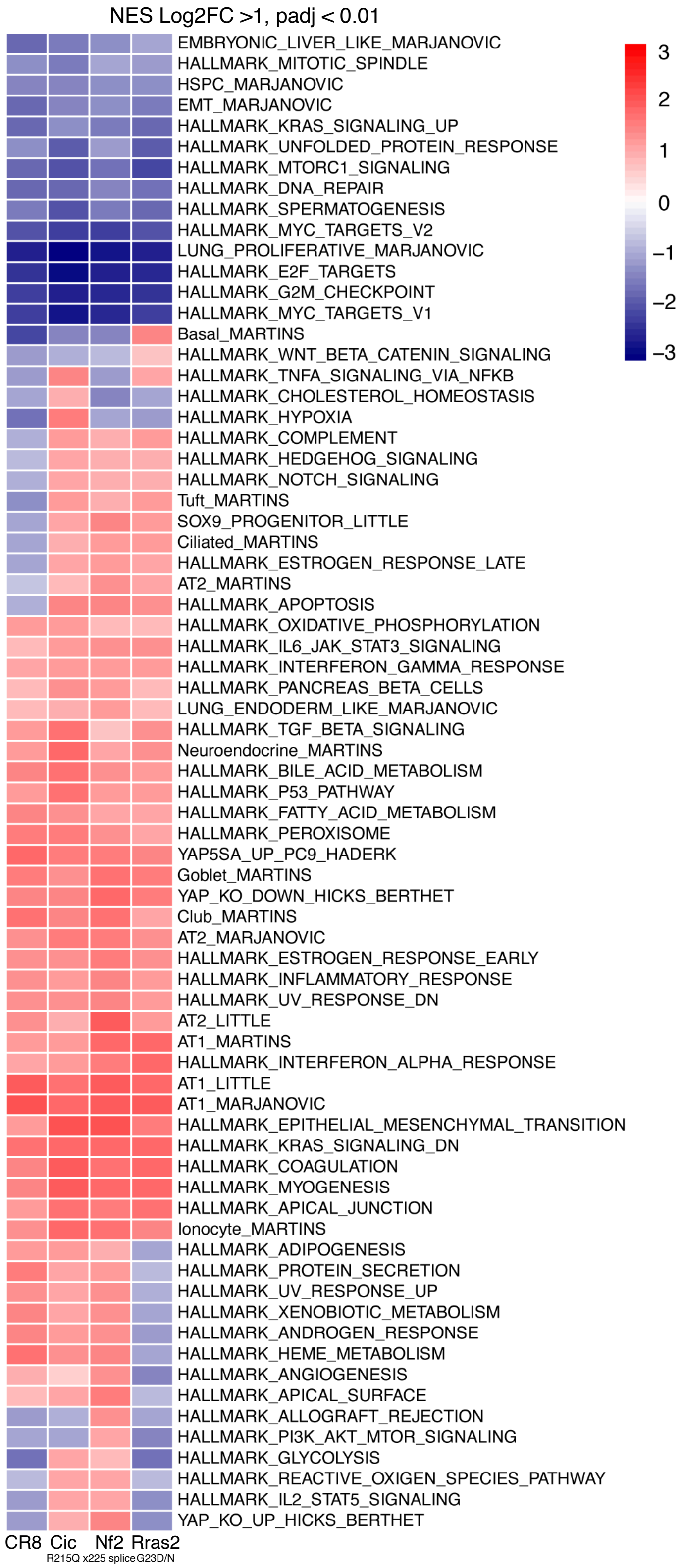


**Supplementary Figure 9. GSEA heatmap**. Heatmap showing gene set enrichment analysis colored by average log2 normalized expression (Log2 FC>1 and p-adj <0.01) of control, Nf2$^{x225}$ splice, Rras2$^{G23D/N}$, and Cic$^{R215Q}$ mutant cells treated with RMC-7977 30nM.
