## Supplementary Methods for "Genetic mechanisms of resistance to targeted KRAS inhibition"

### Supplementary Methods: BEquant

#### Quantifying and profiling sgRNA editing outcomes

Following pre-processing of the paired-end sequencing data to remove low quality reads, we process each read sequentially. Each read consists of two components: *i)* a *sgRNA* and *ii)* a matching *sensor* which records editing. For each read, we perform the following three steps:

1. Extract the sgRNA and sensor from the read by performing an exact string match on the context (scaffold, etc.) which surrounds both the sgRNA and sensor. If the exact match to extract the sgRNA and sensor fails, drop the read.
2. To associate the read with a sgRNA, find the nearest sgRNA in the screen's whitelist (to the (*potentially edited*) sgRNA appearing in the read.
3. Once the associated sgRNA is found, count the number and type of edits at both the read's sgRNA and sensor by comparing it to the associated sgRNA in the whitelist.

The first and third step of the quantification pipeline are straightforward and efficient to implement. Specifically, the first step consists of three exact string matches to extract the sgRNA and sensor, and the third step consists of a single pass through both the read's sgRNA and sensor. In contrast, the second step, which we call *nearest neighbor search* following computer science terminology, is quite challenging to solve efficiently.

To efficiently find the nearest neighbor of the read sgRNA in the whitelist, we developed a probabilistic matching algorithm using the technique of *locality sensitive hashing* [1] popularized in the theoretical computer science literature. However, to draw contrast and benchmark our approach, we first describe a simple, but slow, method for performing nearest neighbor search.

#### Nearest neighbor search using an exhaustive whitelist scan

The simplest approach for finding the nearest sgRNA in the whitelist to the read's sgRNA is as follows. To find the nearest sgRNA, scan (or iterate) through all the sgRNAs in the whitelist and pick the whitelist sgRNA which is closest to the read's sgRNA as quantified using any notion of distance, such as the edit distance. While simple to describe, the approach is quite slow as the time it takes to process each read is proportional to the length of the whitelist. Since the sgRNA whitelist used in our experiments consists of thousands of sgRNAs, this approach is intractable without extensive parallel computing.

#### Nearest neighbor search using locality sensitive hashing

Rather than iterate through all possible sgRNAs in the screen's whitelist for each read as in the preceding approach, we designed a locality sensitive hashing algorithm [1] to efficiently find the nearest neighbor of the read's sgRNA in the whitelist. Using this approach, the time required to process each read becomes much smaller than the size of the whitelist and is tractable even with limited computational resources.

As a warmup, suppose that no editing occurred at the read's sgRNA. Then, the read's sgRNA would exactly match a unique sgRNA in the whitelist and the nearest sgRNA would be itself. In this case, we could find the nearest sgRNA in the whitelist by pre-processing the whitelist into a hash table and looking up the sgRNA in constant time by its hash key. For previous editors, such as FNLS and FNLS-NG, which did not exhibit self-editing, we took exactly this approach [2]. This avoided having to scan through the entire whitelist for each read, but did depend upon the lack of self-editing.

When self-editing does occur, the simple hashing approach no longer works as the read sgRNA does not exactly match the whitelist sgRNA to which it corresponds, leading to different hash keys for the read and whitelist sgRNA. Our proposed fix to resolve this issue is to use what is called a *locality sensitive hash function* [1]. To compute the hash of an sgRNA using locality sensitive hashing, we first randomly mask a fixed subset of nucleotides to obtain a new masked string. Next, we feed the masked string into a standard hash function, such as MD5, to obtain our hash value. The idea is that, if sgRNA's are close together, after randomly masking a subset of nucleotides they become identical with high (e.g. 25%) probability. However, if sgRNA's are quite distinct, even after masking a subset of nucleotides, the strings remain distinct with high probability. Using locality sensitive hashing, similar sgRNAs will hash to the same value and dissimilar sgRNAs will hash to a different value with high probability.

For finding the nearest neighbors to all read sgRNAs, we perform the following steps. First, we fix a random mask to define our locality sensitive hash function, as described previously. Second, we pre-process our whitelist into a hash table by feeding each whitelist sgRNA into the locality sensitive hash function, storing a list of whitelist sgRNAs as values in the hash table. Then, for each read sgRNA, we compute its hash using the locality sensitive hash function and use the hash key to look up the entry in the hash table. We then scan through the associated sgRNAs using the naïve approach and select the closest sgRNA to the read sgRNA. With high probability, the entry of the hash table will contain a whitelist sgRNA that is close to the read, and few whitelist sgRNAs far from the read.

To improve the probability of finding the nearest whitelist sgRNA to a given read sgRNA, rather than store a single table for one locality sensitive hash function, we store multiple tables for a set of random locality sensitive hashing functions. Then, to find the nearest neighbor of a read sgRNA, we look up the read sgRNA in *all* the hash tables to obtain a candidate set of whitelist sgRNAs. We then scan through the candidate set as in the naïve approach, picking the closest whitelist sgRNA. The advantage of using multiple tables is that it provides higher odds of finding the nearest neighbor while trading off with increased runtime.

**A formal description of nearest neighbor search using locality sensitive hashing**

Suppose the set of $m$ screened sgRNAs are provided as a whitelist $\mathcal{W} = \{w_1, \dots, w_m\}$ where $w_i \in \Sigma^{20}$ is a *whitelist sgRNA* and $\Sigma = \{A, T, C, G\}$ refers to the DNA alphabet. Let $\mathcal{R} = \{R_1, \dots, R_n\}$ denote the set of $n$ raw reads in our sequencing experiment. After pre-processing the reads to extract the sgRNA and target site, we obtain a set of $n$ sgRNA-target pairs $\mathcal{T} = \{(g_1, t_1), \dots, (g_n, t_n)\}$ with $g_i \in \Sigma^{20}$ denoting the i$^{th}$ *read sgRNA* and $t_i \in \Sigma^*$ denoting the i$^{th}$ *target site*. We assume that no insertion or deletions (*indels*) occur in the read sgRNA, but that they can occur in the target sites.

The aim of our matching procedure is to find, for each sgRNA-target pair $(g_i, t_i)$ the closest sgRNA $w_j$ in the whitelist. Specifically, we want to find the whitelist sgRNA $w_j$ such that the Hamming distance to the read sgRNA, $H(g_i, w_j)$, is minimized. If several such $w_j$ exist, we arbitrarily pick the first one. The naïve procedure to solve this problem computes $H(w_i, g_j)$ for all pairs of read sgRNAs $g_i$ and whitelist sgRNAs $w_j$. However, since there are $n$ read sgRNAs and $m$ whitelist sgRNAs, the time complexity of this approach would be $\mathcal{O}(nm)$ which is prohibitive in this application.

Instead, we first pre-process the whitelist to efficiently support this matching procedure. To do this, we design $k$ random hash functions $h_1, \dots, h_k$ which *i)* map items of low Hamming distance to the same hash value and *ii)* map items of large Hamming distance to distinct hash values with high probability. Using these $k$ hash functions, we then construct $k$ tables $D_1, \dots, D_k$ by applying the hash function to each whitelist sgRNA, storing the hash values as keys in the table and whitelist sgRNAs as the values. If several whitelist sgRNA's have the same key (i.e. $h_i(g_j) = h_i(g_l)$ for $j \neq l$), we store a set of whitelist sgRNA's as the entry in the table. More formally, entry $x$ in table $D_j$, denoted as $D_j[x]$, stores the set $D_j[x] = \{w \in \mathcal{W} : h_j(w) = x\}$, or the inverse image the hash function $h_j$ at $x$.

Each hash function $h_i$ is defined by a random subset $B_i \subset \{1, \ldots, 20\}$ of nucleotides. Then, to compute $h_i(g)$, the following operations are performed:

1. Construct the string $g' = g[B_i]$ by selecting nucleotides at indices $B_i$.
2. Return the hash value of $g'$ using a standard hashing algorithm.

In our implementation of the probabilistic matching algorithm, we take as user input the size $|B_i|$ and selected $B_i$ by sampling from $\{1, \ldots, 20\}$ without replacement. Hash functions constructed in this way are called *locality sensitive* [1] since two close sgRNAs are likely to have the same hash value. For example, if sgRNAs $g_1$ and $g_2$ differ at only a single nucleotide and $|B_i| = 7$, the probability that they hash to same value, i.e. $h_i(g_1) = h_i(g_2)$, is 0.65.

Once the hash functions $h_i$ and tables $D_i$ have been constructed, which takes a total of $O(nk)$ time, we process each read sgRNA $w_i$ as follows:

1. Construct a set $\mathcal{C}$ of candidate whitelist sgRNAs, which is initially empty.
2. For each of the $k$ hash functions, compute $h_j(w_i)$ and add all whitelist sgRNAs at entry $D_j[h_j(w_i)]$ of the table to $\mathcal{C}$.
3. Compute the Hamming distance $H(c, w_i)$ for all the candidate whitelist sgRNAs in $c \in \mathcal{C}$. Report $c$ with the smallest Hamming distance to $w_i$.

The intuition for the procedure is simple: whitelist sgRNAs $g_j$ at a small Hamming distance from the read sgRNA $w_i$ are likely to end up in the candidate set $\mathcal{C}$ since with high probability they will likely hash to the same key. In contrast, whitelist sgRNAs at a large Hamming distance from the read sgRNA are unlikely to end up in the candidate set, since they will likely hash to distinct keys. Consequently, the candidate set $\mathcal{C}$ is usually small while still containing the closest whitelist sgRNA.

As an example of the effectiveness of our procedure, suppose that a whitelist sgRNA $w$ differs from a read sgRNA $g$ at 3 nucleotides and $|B_i| = 7$. Then, the probability that $h_i(g) = h_i(w)$ for a single hash function $h_i$ is approximately equal to 0.25. However, the odds that $h_i(g) = h_i(w)$ for at least one hash function $h_i$ is approximately equal to $1 - (1 - 0.25)^k = 0.996$ for $k = 20$, implying that the whitelist sgRNA $w$ is almost always in the candidate set $\mathcal{C}$. On the other hand, suppose the whitelist sgRNA $w$ differs from the read sgRNA at 9 nucleotides. Then, the probability that $h_i(g) = h_i(w)$ for a single hash function $h_i$ is approximately equal to $0.004$ and the probability that $h_i(g) = h_i(w)$ for at least one hash function is $1 - (1 - 0.004)^k = 0.08$ for $k = 20$. Consequently, the far away whitelist sgRNA $w$ is excluded from the candidate set $\mathcal{C}$ with high probability.

Increasing the number of positions (k) for the hash key may improve accuracy but also increase false negatives as edited sgRNA reads may be missed and discarded. Conversely, reducing k too far may produce false positives due to high similarity between hash indexes of independent sgRNAs. We tested read assignment across a range of m and k values, identifying k = 20 and m = 7 as a good balance between accurate read assignment, and processing time.

For a full theoretical analysis of the false negative rate (i.e. probability of missing a close sgRNA) and expected runtime (i.e. expected size of the candidate set) for our probabilistic matching procedure, we refer the reader to [1].
